# H4K20me3 controls Ash1-mediated H3K36me3 and transcriptional silencing in facultative heterochromatin

**DOI:** 10.1101/2022.11.25.517763

**Authors:** Mareike Möller, John B. Ridenour, Devin F. Wright, Michael Freitag

## Abstract

Facultative heterochromatin controls development and differentiation in many eukaryotes. In metazoans, plants, and many filamentous fungi, facultative heterochromatin is characterized by transcriptional repression and enrichment with nucleosomes that are trimethylated at histone H3 lysine 27 (H3K27me3). While loss of H3K27me3 results in derepression of transcriptional gene silencing in many species, additional up- and downstream layers of regulation are necessary to mediate control of transcription in chromosome regions enriched with H3K27me3. Here, we investigated the effects of one histone mark on histone H4, namely H4K20me3, in the fungus *Zymoseptoria tritici*, a globally important pathogen of wheat. Deletion of *kmt5*, the gene encoding the sole methyltransferase responsible for H4K20 methylation, resulted in global derepression of transcription, especially in regions of facultative heterochromatin. Reversal of silencing in the absence of H4K20me3 not only affected genes but also a large number of novel, previously undetected, non-coding transcripts generated from regions of facultative heterochromatin on accessory chromosomes. Transcriptional activation in *kmt5* deletion strains was accompanied by a complete loss of Ash1-mediated H3K36me3 and chromatin reorganization affecting H3K27me3 and H3K4me2 distribution in regions of facultative heterochromatin. Strains with a H4K20M mutation in the single histone H4 gene of *Z. tritici* recapitulated these chromatin changes, suggesting that H4K20me3 is essential for Ash1-mediated H3K36me3. The Δ*kmt5* mutants we obtained are more sensitive to genotoxic stressors and both, Δ*kmt5* and Δ*ash1*, showed greatly increased rates of accessory chromosome loss. Taken together, our results provide insights into a novel, and unsuspected, mechanism controlling the assembly and maintenance of facultative heterochromatin.

**Significance:** Facultative heterochromatin contains genes important for specific developmental or life cycle stages. Transcriptional regulation of these genes is influenced by chromatin structure. Here, we report that a little studied histone modification, trimethylation of lysine 20 on histone H4 (H4K20me3), is enriched in facultative heterochromatin and important for transcriptional repression in these regions in an important agricultural pathogen. Furthermore, normal levels of H4K20me3 are essential for deposition of another repressive histone mark, Ash1-mediated H3K36me3, and affect the distribution of other marks including H3K27me3. We conducted the first genome-wide assessment of H4K20 methylation levels in a fungus, and our discoveries reveal that multiple chromatin modifications are required to establish transcriptional silencing, providing the framework to understand epistasis relationships among these histone marks.

## Introduction

Chromatin, the assembly of DNA, RNA, and proteins that constitutes chromosomes, can assume active and inactive states that are correlated with different histone and DNA modifications (1). Transcriptionally inactive, “silent” chromatin is separated into “constitutive heterochromatin”, which is most often marked by H3K9me2/3 and DNA methylation and found within or near centromeres, subtelomeric regions or telomere repeats, rDNA, and transposable elements, and “facultative heterochromatin”, which is typically enriched with H3K27me3 and found on specific genes (2, 3), in larger BLOCs (broad local enrichments) in mouse (4), or across long sections of chromosomes in many fungi (5, 6). Facultative heterochromatin is more dynamic than constitutive heterochromatin, and “on” or “off” states vary between cell types or individuals within a species (7). While H3K27me3, mediated by Polycomb Repressive Complex 2 (PRC2), is considered to be a hallmark histone modification correlated with facultative heterochromatin (8), little is known about other chromatin marks that are important for formation, maintenance, and gene silencing in these regions. In animals, the PRC1 complex and H2AK119ub1 play an important role in PRC2 and H3K27me3 recruitment (9), but so far there is no evidence for a canonical PRC1-like complex in fungi (10).

Other, less studied, histone modifications found in transcriptionally silent regions of facultative heterochromatin include H4K20 and H3K36 methylation. Depending on the organism, one or multiple histone methyltransferases (HMTs) are responsible for methylating H4K20 (11). In animals, one HMT mediates H4K20me1, for example KMT5A in humans and PR-SET-7 in *Drosophila melanogaster* (12, 13), and at least one additional HMT is responsible for H4K20me2/3, for example KMT5B/C in humans and Su(var)4-20 in *D. melanogaster* (14, 15). In contrast, a single enzyme, Kmt5, called Set9 in *Schizosaccharomyces pombe* and SET-10 in *Neurospora crassa*, catalyzes all H4K20me in fungi (16–18). Presence of the different methylation states of H4K20 (i.e., me1, me2, or me3) are essential for development and associated with DNA repair, replication, cell cycle control, genome stability, and chromatin compaction in animals (11, 19–24). In contrast, there are few studies on fungi addressing the function of H4K20me. In fission yeast, H4K20me is important for DNA repair by recruitment of Crb2 to sites of DNA damage (16). Studies with the filamentous plant pathogenic fungi *Fusarium graminearum*, *F. fujikuroi*, and *Magnaporthe oryzae* (syn. *Pyricularia oryzae*) revealed minor, if any, effects on virulence (17, 18), and only subtle and species-specific effects on expression of secondary metabolite cluster genes and tolerance to stress conditions in Δ*kmt5* strains (17). Surprisingly, none of the previous studies analyzed the genome-wide distribution of any of the H4K20me modification states in fungi.

In contrast, H3K36me is a widespread and well-studied histone modification deposited by Set2 in an RNAPII-dependent manner during transcript elongation (25) and, in some organisms, by another group of RNAPII-independent SET-domain proteins, the ASH1 or NSD family proteins (26, 27). ASH1-mediated H3K36me is linked to Polycomb-mediated repression and is thought to counteract PRC2-mediated H3K27me3 (28, 29). In fungi, ASH1-mediated H3K36me3 co-occurs in regions enriched with H3K27me3 and is important for normal distribution of this mark, at least in *N. crassa* and *F. fujikuroi* (30, 31).

In this study, we wished to address how H4K20me and ASH1-mediated H3K36me3 contribute to the formation, maintenance, and silencing of facultative heterochromatin. The filamentous fungus *Zymoseptoria tritici*, a global pathogen of wheat, has a well-characterized epigenetic landscape, which allowed us to differentiate euchromatin, constitutive or facultative heterochromatin, regions with cytosine methylation (5mC), short regional centromeres, subtelomeric regions, and telomere repeats (6, 32, 33). Facultative heterochromatin, enriched with H3K27me3 and transcriptionally silenced, covers subtelomeric regions and accessory chromosomes (also referred to as “non-essential” or “conditionally dispensable” chromosomes) that create variation between isolates (6, 34). While H3K27me3 in these regions is involved in chromosome stability and accumulation of mutations, loss of H3K27me3 has only minor impacts on transcriptional activation (32, 35). Therefore, we hypothesized that another repressive histone modification exists that is epistatic to H3K27me in *Z. tritici*, and perhaps other organisms.

Here we show that H4K20me3 is one such modification and that it is generated exclusively by the SET-domain HMT, Kmt5. Kmt5 and H4K20me3 are crucial for transcriptional silencing in regions of facultative heterochromatin and essential for Ash1-mediated H3K36me3, thereby contributing significantly to formation of H3K27me3-enriched facultative heterochromatin.

## Results

### Kmt5 and Ash1 are important for normal growth but not essential in *Zymoseptoria*

We identified *kmt5* (Zt_chr_3_00475) and *ash1* (Zt_chr_13_00232) in the genome of the reference isolate IPO323 by BLAST searches with *S. pombe* SET9 and *N. crassa* ASH-1 sequences as baits. We found a single H4K20 methyltransferase homolog, consistent with findings in other fungi but different to metazoans, where multiple enzymes are involved in catalyzing H4K20me1 and H4K20me2/3 (Figure 1A). Protein sequence alignments of the SET domains of H4K20 methyltransferases in *Z. tritici*, *S. pombe*, *D. melanogaster, Danio rerio*, and human revealed higher sequence similarity between Kmt5 in fungi and known H4K20me2/3 methyltransferases than H4K20me1 methyltransferases (Figure S1). We showed that Kmt5 in *Z. tritici* is responsible for all H4K20 methylation, as *kmt5* deletion strains are unable to generate either H4K20me1 or H4K20me3 (Figures 1B, S2).

**Figure 1.**
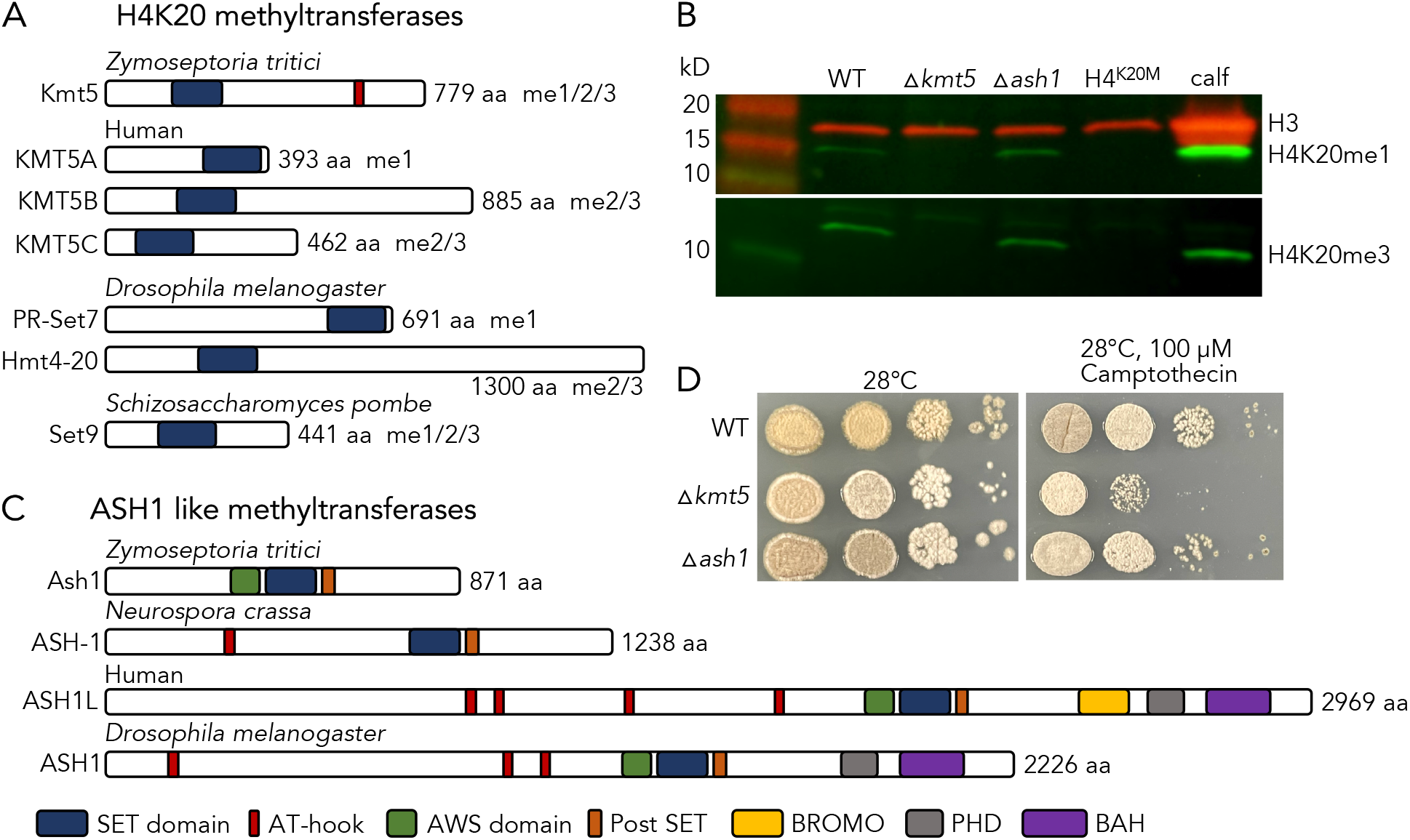
Domain structure and characterization of defects related to two histone methyltransferases involved in generation of facultative heterochromatin. **A)** Comparison of H4K20 methyltransferases related to Kmt5 (accession numbers in Figure S1B). The blue box denotes the SET domain. **B)** Deletion of *kmt5* and substitution of H4 lysine 20 with methionine (H4K20M) causes complete loss of H4K20 methylation but *ash1* deletion has no effect on H4K20 methylation. Calf thymus histones were used as controls. **C)** Comparison of H3K36 Ash1 methyltransferases. **D)** *kmt5* but not *ash1* mutants are mildly affected by the DNA replication inhibitor camptothecin at high temperature.

There is a single gene encoding a *Z. tritici* Ash1 homologue. This Ash1 protein lacks, as does *Neurospora* ASH-1 (30), the C-terminal BROMO, PHD and BAH domains found in animal Ash1 homologues but contains the conserved AWS, SET and post-SET domains (Figure 1C). While the specific functions of the additional domains in most animal Ash1 proteins are not known, the PHD and BAH domain are essential for Ash1 to counteract Polycomb-mediated repression in *D. melanogaster* (36). Absence of these domains in fungal Ash1 homologs suggests different mechanisms of interaction with chromatin and PRC2, specifically.

To determine the function of *kmt5* and *ash1*, we deleted the respective genes in the wild-type isolate Zt09 (37) by replacement with a gene encoding hygromycin phosphotransferase (*hph*), conferring resistance to hygromycin B. Mutant strains were confirmed by PCR and Southern assays (Figure S3). Both deletion mutants exhibit phenotypic differences compared to the wild-type strain. The Δ*kmt5* mutant showed increased sensitivity to genotoxic stress and altered morphology at higher temperatures (28°C *versus* the usual 18°C; Figure 1D). Deletion of *ash1* only slightly retards growth and had overall minor effects on growth under the tested conditions (Figures 1D, S4).

### Loss of Kmt5 and Ash1 result in increased chromosome loss

Previously, we measured accessory chromosome loss as an indicator for overall genome stability (32). A four-week growth experiment including five replicate cultures of wild-type, Δ*kmt5*, and Δ*ash1* strains revealed an increase in accessory chromosome loss in both mutants (Table 1). Increased chromosome loss rates did not affect all accessory chromosomes equally; chromosomes 14 and 16 in Δ*kmt5*, and chromosome 16 in Δ*ash1* were lost much more frequently than any other accessory chromosomes. This indicates that both Kmt5 and Ash1 are important for the maintenance of at least certain accessory chromosomes, and more likely general genome stability.

**Table 1:**
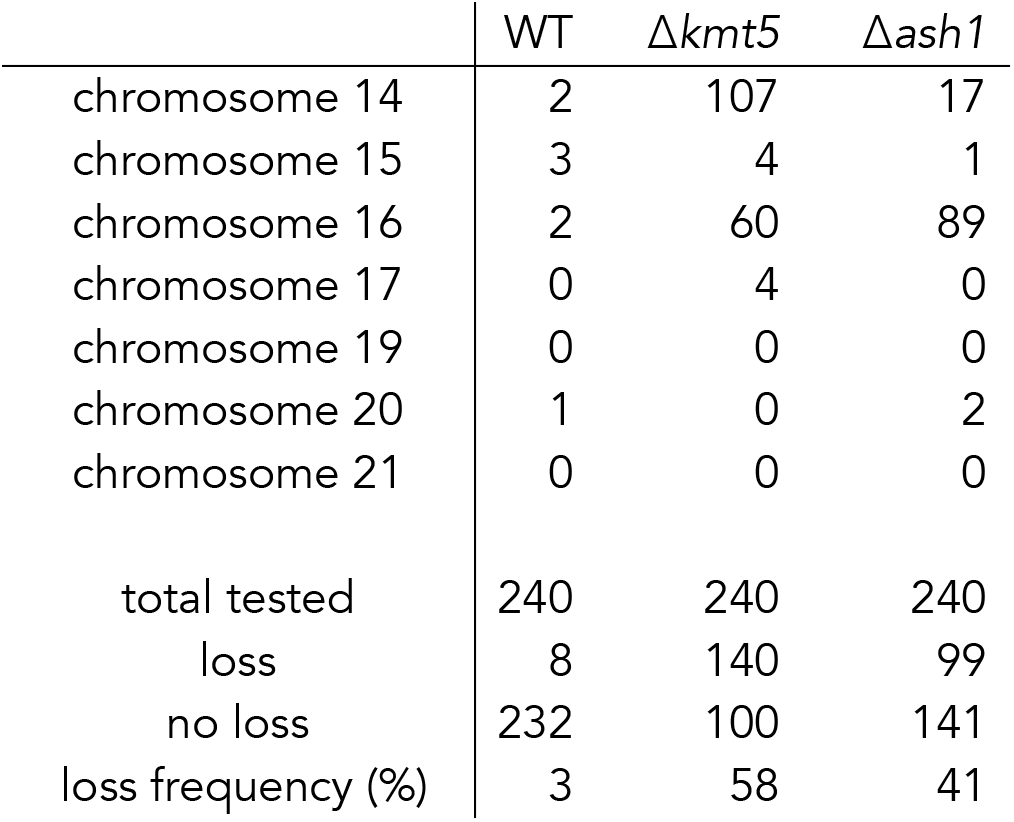
Chromosome loss in wild-type, Δ*kmt5* and Δ*ash1* strains. Chromosome loss is tabulated by individual chromosome and as the sum of all losses (loss), compared to strains without chromosome loss (no loss). Overall chromosome loss frequency (% loss) was calculated as (loss/total tested) *100. Some strains lost multiple chromosomes.

### ChIP-seq uncovers different heterochromatin states in *Z. tritici*

To determine the effects on chromatin in both Δ*kmt5* and Δ*ash1* mutants, we first performed ChIP-seq of selected histone modifications associated with eu- and heterochromatin in wild-type strains. We found that H4K20me3 is widespread along chromosomes, enriched across gene bodies, and exhibits pronounced enrichment in regions we previously characterized as facultative heterochromatin, based on low levels of gene expression and the presence of H3K27me3 (6, 32, 33) (Figures 2A, S5). For the first time, we tested localization of H3K27me2 in *Z. tritici* and found that it is present in the same regions as H3K27me3 but at lower levels of enrichment. H4K20me1 did not show any specific enrichment but was reduced in regions with strong H4K20me3 enrichment, e.g., facultative heterochromatin. H3K36me3 showed a similar pattern and distribution as H4K20me3, and both H3K36me3 and H4K20me3 were absent from constitutive heterochromatin that is marked by H3K9me3 alone, or by a combination of H3K9me3 and H3K27me2/3 (Figures 2A, S5A). To characterize modifications enriched in facultative heterochromatin regions in more detail, we plotted enrichment of H4K20me3 and H3K36me3 within 500 bp windows of regions that are enriched with H3K27me3 in wild type (Figure 2B). We found two distinct clusters within H3K27me3-enriched regions: cluster 1 was enriched with H4K20me3 and H3K36me3, while cluster 2 exhibited very low levels of H3K36me3 and H4K20me3. Further analyses of the genomic context of these clusters revealed that cluster 1 mostly overlaps with genes and intergenic regions, while cluster 2 overlaps with transposable elements (TEs) that are also enriched with H3K9me3, few genes, and intergenic regions (Figure 2C). H3K27me3 and H3K9me3 co-occur in regions close to the telomeres, some repeats, and sections of accessory chromosomes; the common feature of these regions are TEs. Both H3K36me3 and H4K20me3 are predominantly excluded from these regions.

**Figure 2:**
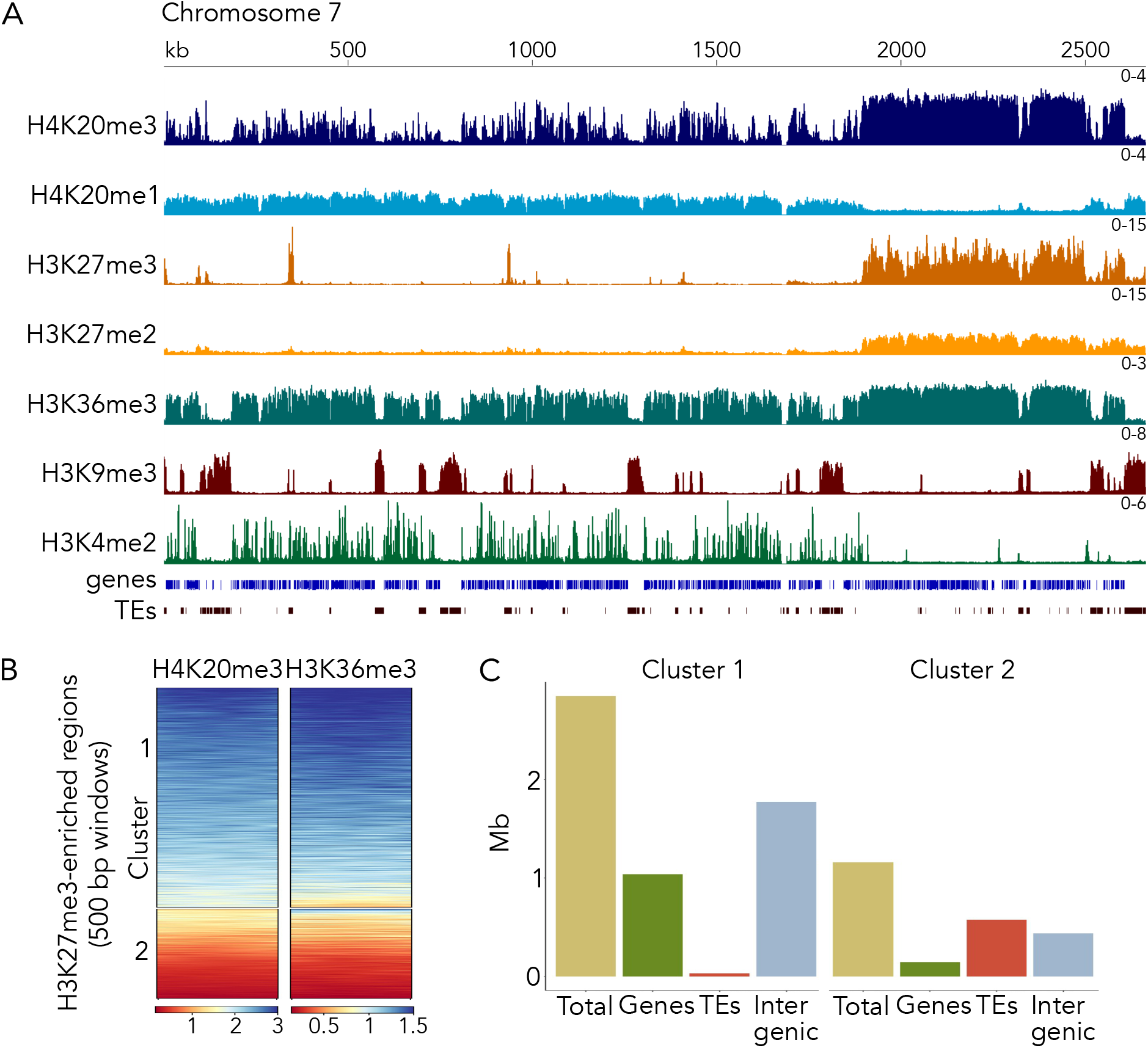
Characterization of different histone modification patterns and their genomic context. **A)** Distribution of selected histone modifications on chromosome 7 in the wild-type *Zymoseptoria tritici* strain. In facultative heterochromatin regions, H4K20me3, H3K27me2/3 and H3K36me3 overlap. **B)** Enrichment of H3K36me3 and H4K20me3 within H3K27me3 regions. There are two distinct clusters, regions where H3K36me3 and H4K20me3 co-occur (cluster 1) and regions where both marks are absent or only one of them is present (cluster 2). **C)** Genomic context of cluster 1 and 2; Mb, million base pairs.

These two clusters may signify functionally different types of facultative heterochromatin, with cluster 1 representing the traditionally accepted type of facultative heterochromatin conditionally repressing genes, while cluster 2 represents constitutive heterochromatin doubly marked with H3K9me3 and H3K27me3. Based on these results we now differentiate at least three types of heterochromatin in *Z. tritici*: (1) constitutive heterochromatin, correlated with H3K9me3 enrichment only and enriched on repeats and TEs, (2) constitutive heterochromatin, correlated with enrichment of H3K9me3 and H3K27me3, found close to telomeres and some repeats and TEs, especially on accessory chromosomes, and (3) facultative heterochromatin, correlated with H3K27me3, H3K36me3, and H4K20me3 enrichment, mostly found on genes and intergenic regions either in large blocks or in shorter segments dispersed on chromosomes. The functional relevance of these states remained to be determined and here we began these investigations.

### Facultative heterochromatin marked by H4K20me3, H3K36me3, and H3K27me3 shows strong interactions

To further characterize chromatin states and their influence on spatial interactions in the nucleus, we performed Hi-C experiments to find regions in the genome that physically interact. We compared wild type and a Δ*kmt6* mutant to see if loss of H3K27me3, the hallmark modification of facultative heterochromatin, impacts interaction frequencies in specific regions. We found strong interactions between regions enriched with H3K27me3 that we characterized as facultative heterochromatin (cluster 1) in wild type (Figure 3). As in other organisms (38–41) centromeres, which cannot be identified by a single chromatin mark in *Z. tritici* (6), interact most strongly. Large blocks of facultative heterochromatin are found on accessory chromosomes and on the right arm of chromosome 7. These blocks show strong inter- and intrachromosomal interactions but few interactions with non-facultative heterochromatin regions (Figure 3), revealing that they are physically separated from euchromatin. When comparing results from wild type to those from the Δ*kmt6* mutant, we found that these strong interactions between facultative heterochromatin regions persist, even in absence of H3K27me3 indicating that other mechanisms and not H3K27me3 alone are responsible for these strong interactions. Notably, H4K20me3 enrichment was not affected by absence of H3K27me3 in the Δ*kmt6* mutant (Figures 3B, S6).

**Figure 3:**
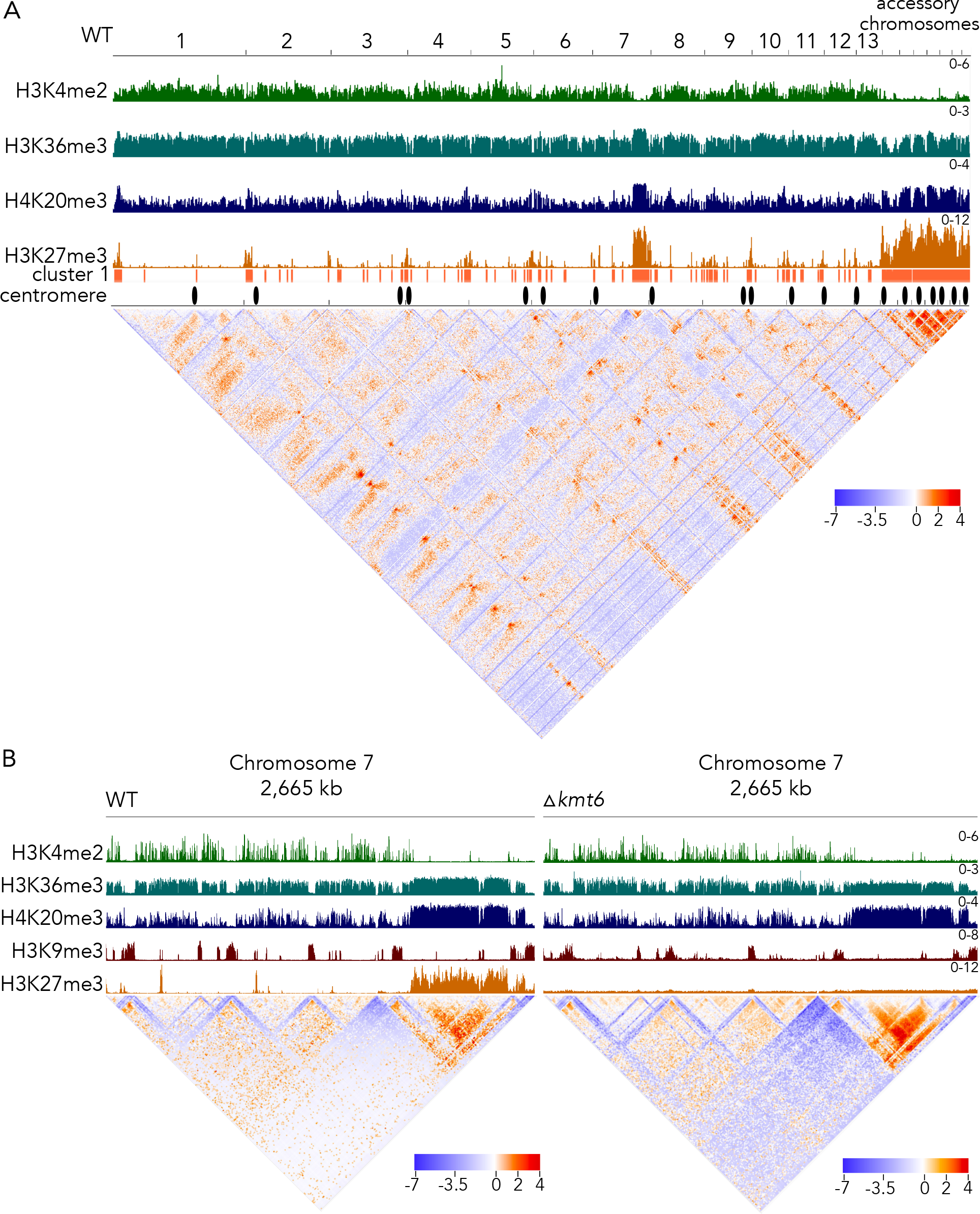
Genome-wide and chromosome-specific Hi-C interaction map in wild-type and Δkmt6 mutant. **A)** Interacting genomic regions in the wild-type genome. Shown are log_2_ ratios of observed interactions relative to expected interactions; fewer interactions than expected are colored in blue and more interactions than expected are colored in red. Most centromeres (6) and regions of facultative heterochromatin (cluster 1) show strong interactions. **B)** Side-by-side comparison of interactions on chromosome 7 in wild-type and the Δ*kmt6* mutant. The right arm of chromosome 7 shows strong regional interactions but very few interactions with the left arm of the chromosome. The interacting region correlates with H3K27me3 enrichment in WT, but interactions do not change in absence of H3K27me3.

### Kmt5 is essential for Ash1-mediated catalysis of H3K36me3 and normal H3K27me3 distribution

Deletion of *kmt5* resulted in complete loss of H4K20me3 (Figures 1B and 4) and impacted the distribution of several other histone modifications tested (Figures 4 and S7, S8). Most prominently, we found that H3K36me3 was lost from most regions in which it co-localized with H3K27me3 in wild type. The same regions lost H3K36me3 in the *ash1* deletion mutant, indicating that Ash1 activity is compromised in the *kmt5* mutant. Loss of Ash1, however, does not impact H4K20me3, indicating that H4K20me3 is epistatic to, or acts upstream of, Ash1-mediated H3K36me3.

**Figure 4:**
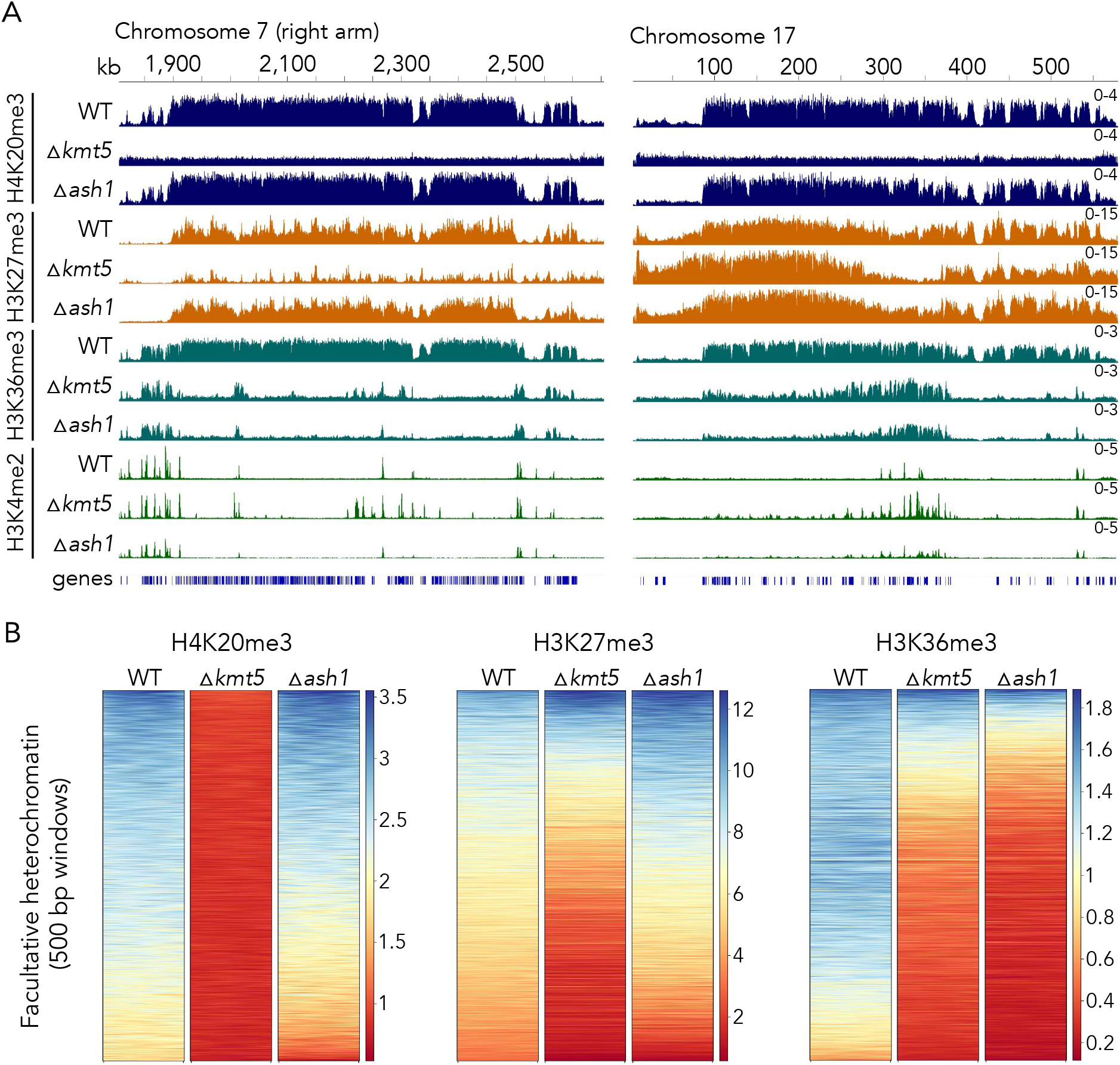
Effects of deletion of *kmt5* or *ash1*, encoding two histone methyltransferases involved in the regulation of H4K20me3 and H3K36me3 levels. **A)** In the absence of Kmt5, all H4K20 methylation is removed but H4K20me3 enrichment is not affected by lack of Ash1. Lack of Kmt5 or Ash1 alters H3K27me3 in selected regions of the genome, and the lack of either Kmt5 or Ash1 removes H3K36me3 from regions that are not methylated by a second H3K36 methyltransferase, Set2. H3K36 methylation by Set2 is largely correlated with H3K4me2. These results strongly suggest that Kmt5 controls at least some H3K36 methylation, which in turn controls H3K27me3 in some regions of the genome. **B)** Comparison of enrichment of histone modifications in regions of facultative heterochromatin. Deletion of *kmt5* results in loss of H4K20me3, overall reduction in H3K27me3, and strong reduction in H3K36me3. Deletion of *ash1* affects H4K20me3 and H3K27me3 only mildly but strongly reduces H3K36me3 in these regions.

While impacts on H3K36me3 were similar in *kmt5* or *ash1* deletion strains, loss of *kmt5* had more severe effects on H3K27me3 enrichment. In the absence of Kmt5, we observed major changes in H3K27me3 distribution, including losses of large blocks of H3K27me3, some movement to constitutive heterochromatin, and losses and gains of peaks in numerous shorter dispersed peaks. In addition, loss of H4K20me3, H3K36me3, and H3K27me3 in the Δ*kmt5* mutant was often accompanied by gain of H3K4me2, suggesting a transition from facultative heterochromatin to transcriptionally active euchromatin (Figure 4A). In contrast, we detected fewer differences in H3K27me3 enrichment in the *ash1* mutant, mostly losses or gains in a limited number of the shorter dispersed peaks and increased enrichment in large H3K27me3-enriched blocks on accessory chromosomes.

We found that most changes in chromatin states in Δ*kmt5* and Δ*ash1* mutants occurred in regions that we characterized as facultative heterochromatin (cluster 1), i.e., regions that show enrichment for H3K27me3, H3K36me3, and H4K20me3, but to a much lesser extent in regions outside of facultative heterochromatin (Figure S7). Looking specifically at enrichment of the three marks in those regions in wild type and mutants (Figure 4B), we found that, as expected, H4K20me3 is absent in Δ*kmt5* but only slightly reduced in Δ*ash1*. H3K27me3 was reduced in many regions in Δ*kmt5* but slightly increased in Δ*ash1*, and H3K36me3 was greatly reduced in Δ*ash1* and only slightly less so in Δ*kmt5* when compared to enrichment in wild type. Taken together, we showed that Kmt5 is important for facultative heterochromatin assembly and maintenance and that it is essential for Ash1 activity.

### H4K20M mutation alters abundance and distribution of H4K20me3

To test whether the presence of the Kmt5 protein or its catalytic activity, i.e., methylation of H4K20, is important for H3K36me3, we constructed histone H4K20 mutants, replacing lysine residue 20 with a methionine. We generated two different types of mutants, one a clean replacement of the single endogenous histone H4 gene (*hH4*) by *hH4^K20M^* (*H4^K20M^*) and the second an ectopic addition of *hH4^K20M^*, generating the potential for a mixed population of nucleosomes (*H4^K20/K20M^*). The replacement allele recapitulated results obtained with the Δ*kmt5* mutant, showing complete absence of H4K20me3, loss of Ash1-mediated H3K36me3, and reduced H3K27me3 enrichment, revealing that presence of the histone mark, but not necessarily presence of the Kmt5 protein, is essential for Ash1 recruitment (Figures 5A-C, S8). In contrast, we observed striking differences in H4K20me3 distribution in the strain with the ectopic *hH4^K20M^* integration (*H4^K20/K20M^*). In this mutant, H4K20me3 enrichment was retained only in specific regions, co-localizing with Ash1-mediated H3K36me3 and H3K27me3 but lacking from the rest of the genome. Analysis of the regions enriched with H4K20me3 in the *hH4^K20/K20M^* mutant revealed that H3K36me3 was still slightly reduced compared to wild type, but levels were higher than in both Δ*kmt5* and *hH4^K20M^* mutants, while H3K27me3 enrichment was comparable to wild type (Figure 5A-C). Presence of H4K20me3 in these regions suggests that these regions are preferential targets of Kmt5. Based on data collected with mutant histone H4 alleles, presence of H4K20me3 appears to be necessary to deposit H3K36me3 and H3K27me3 and thus for maintenance of a facultative heterochromatin state.

**Figure 5:**
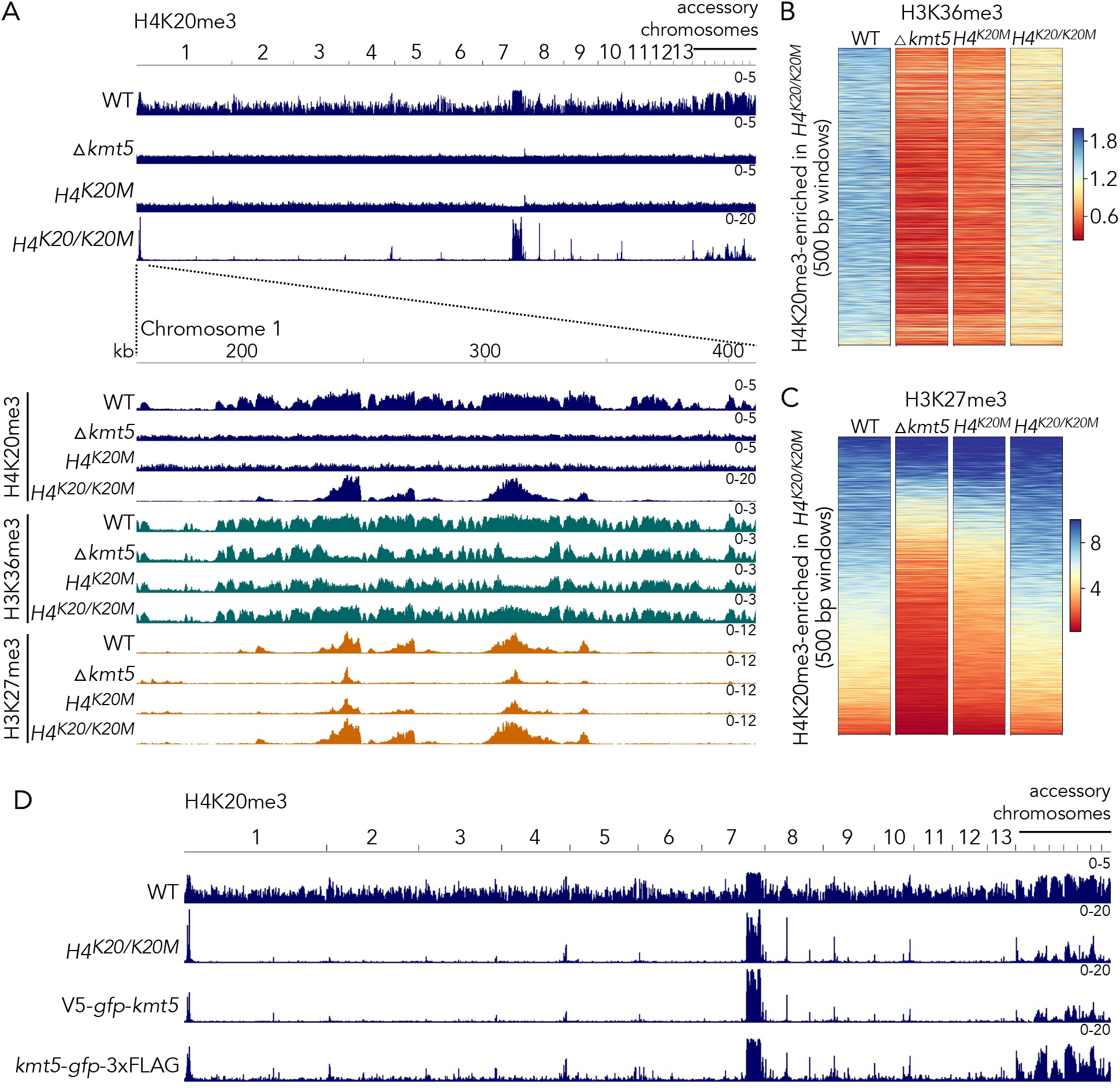
An H4^K20M^ substitution phenocopies deletion of *kmt5* but heterozygous H4^K20/K20M^ strains show specific H4K20me3 in regions of facultative heterochromatin in wild type. **A)** H4K20me3 distribution in Δ*kmt5* and histone mutants. **B)** and **C)** Effects of deletion of *kmt5* or substitution to H4^K20M^ and H4^K20/K20M^ on regions that retain H4K20me3 in H4^K20/K20M^ strains. **B)** H3K36me3 is severely reduced in Δ*kmt5* and H4^K20M^ but partly restored in H4^K20/K20M^. **C)** H3K27me3 is reduced in *kmt5* and H4^K20M^ but similar to wild-type levels in H4^K20/K20M^. **D)** A similar effect on H4K20me3 observed with heterozygous H4^K20/K20M^ is found when Kmt5 is tagged with GFP on either the amino or carboxy terminus.

### Tagging of Kmt5 impairs protein function and limits H4K20me3 spreading

In order to characterize Kmt5 in more detail, we generated strains harboring N-terminally (*V5*-*gfp*-*kmt5*) or C-terminally (*kmt5*-*gfp*-*3xFLAG*) tagged Kmt5 alleles by either replacing the endogenous *kmt5* in wild type or complementing the Δ*kmt5* mutant at the original locus. Based on ChIP-seq analyses, none of these mutants retained or restored wild-type levels of H4K20me3 enrichment. H4K20me3 levels were similar to those observed in the *H4^K20/K20M^* strain in specific locations co-localizing with Ash1-mediated H3K36me3 and H3K27me3 but mostly absent from the rest of the genome (Figure 5D). The C-terminally tagged Kmt5 mutant maintained some H4K20me3 outside of facultative heterochromatin but not at levels comparable to those in wild type.

Notably, integration of the C-terminally tagged Kmt5 in either wild-type or Δ*kmt5* backgrounds led to similar defects in H4K20me3 distribution. This suggests the difference in H4K20me3 is not a result of impaired *de novo* deposition of H4K20me3 after complete loss of the modification but rather a defect in H4K20me3 genome-wide deposition or maintenance associated with impaired function of the tagged Kmt5 protein (Figure S8).

### Eaf3 is essential for Ash1 activity

To further investigate the connection between H4K20me3 and H3K36me3, we searched the genome for predicted proteins that, in other organisms, have been shown to bind to H4K20me3 or to interact with either Ash1 or Kmt5. In *S. pombe*, a PWWP domain-containing protein (Pdp1) occurs in a complex with Kmt5 and is required for all H4K20me (42). We found a single protein with a PWWP domain in *Z. tritici* (Zt_chr_4_873) and other filamentous fungi (Figure 6A). Deletion of the respective gene in *Z. tritici* did not affect H4K20me3 or H3K36me3 (Figure 6B and C) but rather led to increased enrichment of H3K27me3 in numerous regions along the genome (Figures S7, S8, S9), suggesting that there is no homolog of the *S. pombe* Pdp1 in filamentous fungi but that this protein is important for regulation of other chromatin states. A homolog of *Z. tritici* Pdp1 was found to interact with a chromatin remodeling factor, ISWI, in *N. crassa*, where it also increased H3K27me3 levels; thus *Z. tritici* Pdp1 is a putative homolog of *S. cerevisiae* Ioc4 (43).

**Figure 6:**
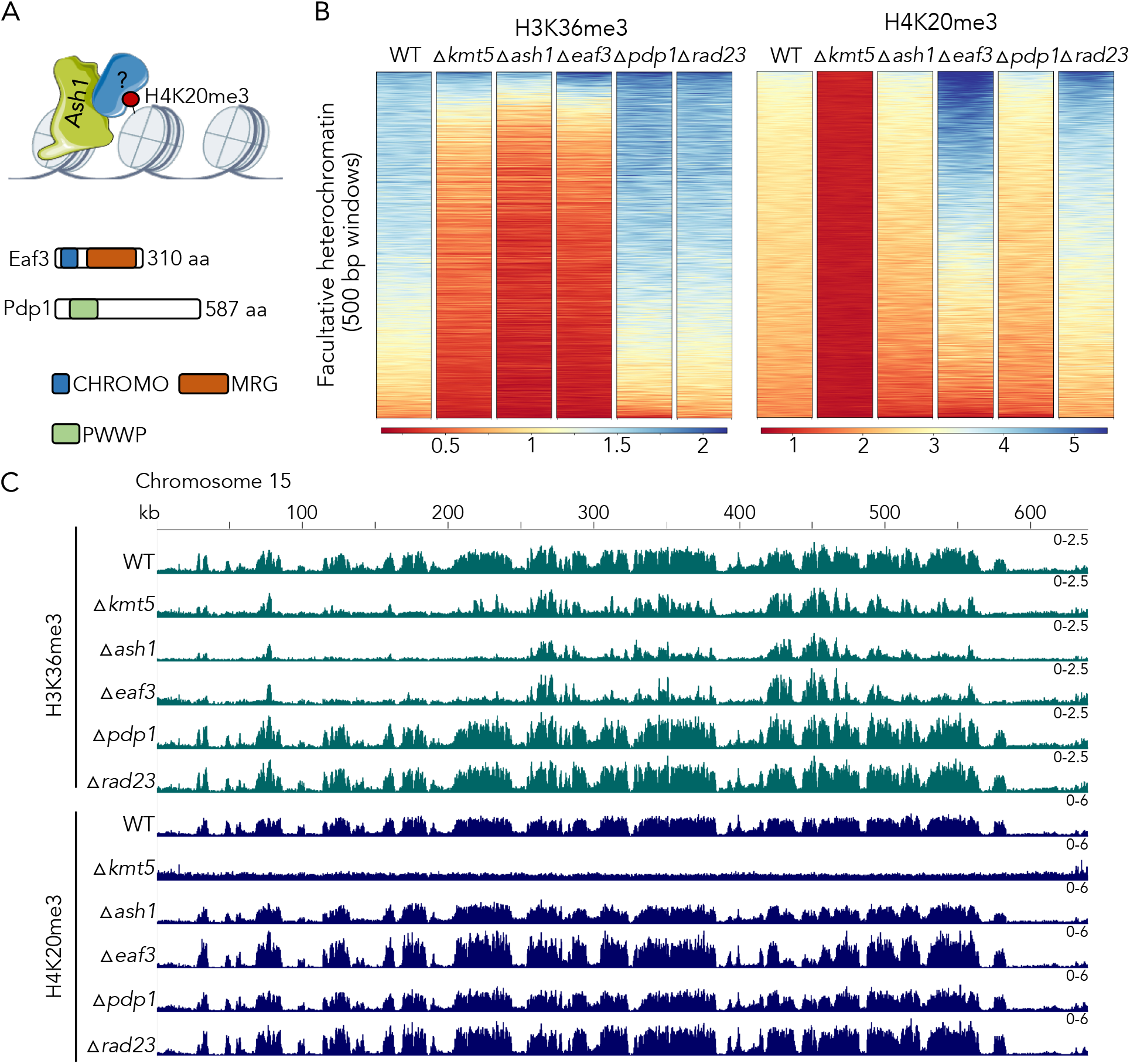
Eaf3 links H4K20 and H3K36 methylation. **A)** A putative H4K20me3-binding protein (“?”) attracts the Ash1 H3K36 methyltransferase, Ash1, to H4K20me3-enriched regions. The PWWP motif protein, Pdp1, is essential for H4K20me in fission yeast, while the CHROMO and MRG domain protein, Eaf3, had been shown to bind H3K36me3 in multiple organisms. **B)** Enrichment of H4K20me3 and H3K36me3 in facultative heterochromatin. We excluded the putative Pdp1 homolog (*pdp1*) because deletion of this gene had no effects on H4K20 or H3K36 methylation when compared to WT. The homologue of Eaf3, however, resulted in a large increase in H4K20me3, and a loss in H3K36me3, suggesting that it may interact with both H4K20me3 and H3K36me3. Loss of the potential H4K20me3 demethylase, Rad23, increased H4K20me3 levels. **C)** ChIP-seq tracks showing H3K36me3 and H4K20me3 on accessory chromosome 15 in WT and Δ*kmt5*, Δ*ash1*, Δ*eaf3*, Δ*pdp1* and Δ*rad23* strains.

Our second candidate protein for connecting H4K20 methylation and Ash1 is a homolog of the chromodomain-containing protein Eaf3 in *S. cerevisiae* or MRG15 in *D. melanogaster*. In yeast, Eaf3 is part of the histone deacetylase complex Rpd3S (44) and the histone acetyltransferase complex NuA4 (45), and its chromodomain can bind to both H3K36me3 and H4K20me3 *in vitro* (46), though not at the same time. *Drosophila* MRG15 interacts with Ash1 and is essential for Ash1 function (47). We deleted the *Z. tritici* homolog of the *eaf3* gene (Zt_chr_1_1455) and performed ChIP-seq on the mutant. Absence of *eaf3* resulted in loss of H3K36me3 in the same regions where we observed loss of H3K36me3 in Δ*ash1* and Δ*kmt5* mutants indicating that Eaf3 is required for Ash1 activity (Figures 6B and C, S7, S8). Eaf3 was not required for H4K20me3; on the contrary, we observed increased levels of H4K20me3 in the Δ*eaf3* mutant in regions that are targeted by Ash1 in wild type (Figure 6B and C). We did not observe a similar increase outside of regions of facultative heterochromatin. Because we did not detect a similar increase in H4K20me3 in Δ*ash1* mutants, it is not absence of H3K36me3 but a mechanism upstream of Ash1 recruitment that influences H4K20me3 levels in Δ*eaf3*.

### A Rad23 homolog may act as a histone H4K20me3 demethylase

A recent study found that two human hHR23 proteins, homologs of *S. cerevisiae* Rad23, act as demethylases of H4K20me1/2/3 in human cells (48). We found one potential homolog of yeast Rad23 in *Z. tritici* (previously not annotated, on chr 2: 3,656,371-3,657,715) and deleted the putative *rad23* gene to determine whether Rad23 is involved in H4K20me3 demethylation in *Z. tritici*. We found increased levels of H4K20me3 in facultative heterochromatin in Δ*rad23* strains and minor differences in H4K20me3 levels outside of these regions indicating that Rad23 may be involved in demethylation of H4K20, particularly in facultative heterochromatin regions (Figure 6B and C, S7, S8).

### H4K20me is required for transcriptional repression in facultative heterochromatin

To investigate whether the observed changes in chromatin structure correlate to changes in gene expression, we sequenced mRNA from three replicates of wild type, Δ*kmt5*, Δ*ash1*, Δ*eaf3*, *hH4^K20M^*, and *hH4^K20/K20M^* strains and determined which genes were differentially expressed (Figures 7A, S10). As a control for the H4^K20/K20M^ strain, we also included a strain that carries an ectopic wild-type H4 copy. We detected few (only 10) differentially expressed genes between wild type and this strain, concluding that an additional wild-type H4 copy has negligible impact under these conditions (Table S1). We showed previously (32) that loss of H3K27me3 alone has surprisingly minor effects on transcription, as only a few genes, mostly located on accessory chromosomes, showed increased levels of expression. In contrast, deletion of *kmt5* results in upregulation of a large set of genes, especially in regions of facultative heterochromatin that are usually enriched with H3K27me3, Ash1-mediated H3K36me3, and H4K20me3. Out of 718 upregulated genes (log_2_fc >| 0.585 |, padj <0.01, Figure 7A) in Δ*kmt5*, 286 are located in facultative heterochromatin (~40% of upregulated genes). Similarly, 39% of upregulated genes in *hH4^K20M^* are within facultative heterochromatin, indicating that Kmt5 and H4K20me3 are especially important for gene repression in those regions. While we also observed upregulation of genes in facultative heterochromatin in the other mutants, their relative proportion is considerably lower (26% in Δ*ash1*, 19% in *hH4^K20/K20M^*, and 15% in Δ*eaf3*). H4K20me3 levels in these mutants were largely unchanged or even increased (Δ*eaf3*) suggesting that presence of H4K20me3 may limit derepression. We found that more than 55% of upregulated genes were shared in Δ*kmt5* and Δ*ash1* mutants and almost 90% of upregulated genes in the *hH4^K20M^* mutant were also upregulated in Δ*kmt5*, Δ*ash1*, or both. In general, a large proportion of upregulated genes are shared in the mutants we sequenced, indicating that they affect a common silencing pathway (Figure S10). Most of the upregulated genes (~94%), as well as genes in facultative heterochromatin in general, lack functional annotations (only 57 out of 976 [~6%] genes have been previously annotated in these regions), making it difficult to assess enrichment for putative functions among those genes. A list including all genes, expression levels, and predicted function can be found in Table S1.

**Figure 7:**
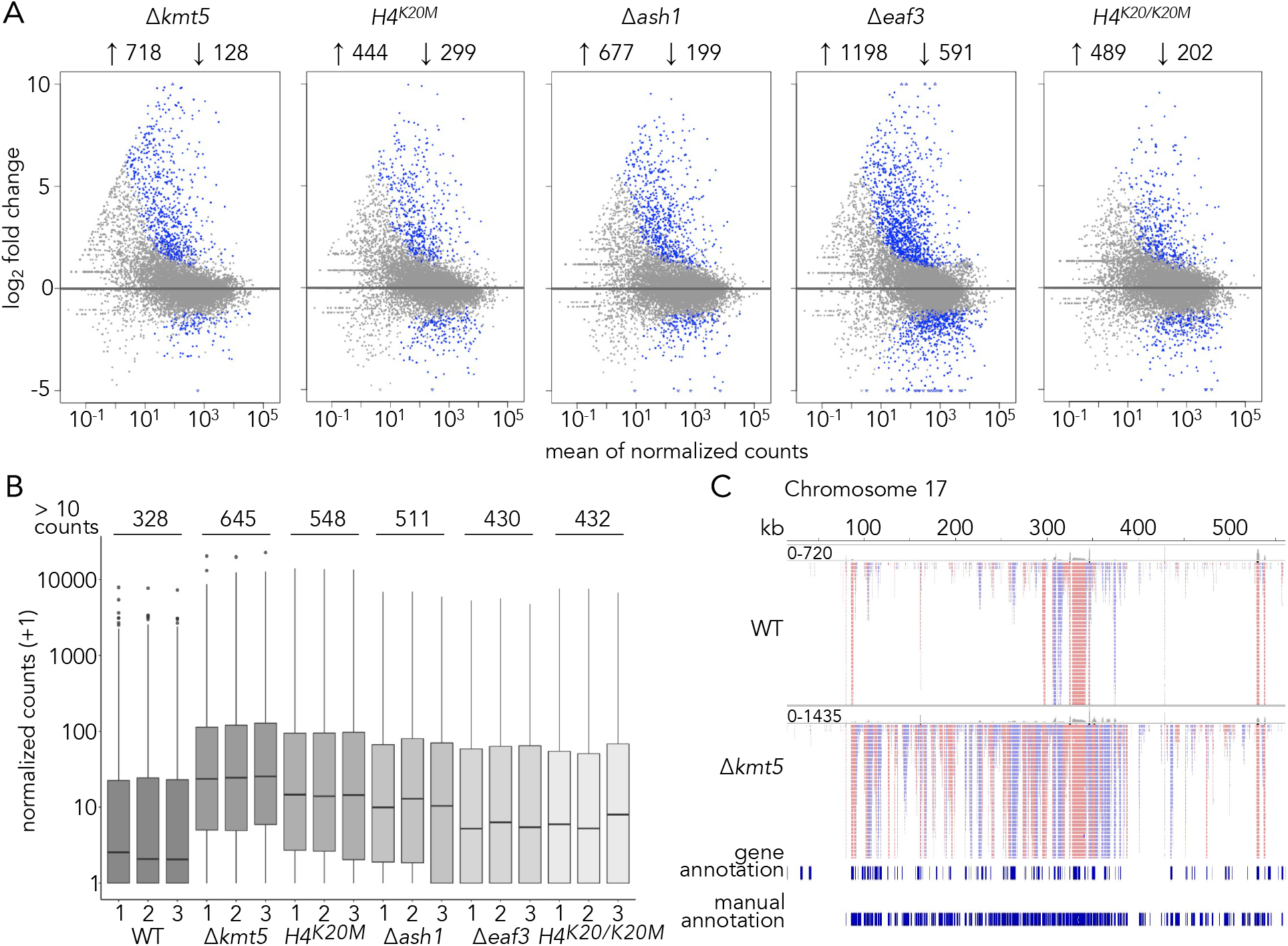
H4K20 methylation is important for transcriptional silencing in facultative heterochromatin. **A)** Results from differential expression analysis (log2fc > | 0.585 |, *padj* < 0.01) comparing wild-type and mutant strains. **B)** Normalized counts assigned to transcripts in facultative heterochromatin; transcripts were newly annotated with stringtie. RNA-seq results on *kmt5, eaf3*, and histone mutants, as well as on wild-type (WT) samples revealed a large group of novel transcripts, especially on accessory chromosomes, that had not been detected before and that come from H3K27me3-enriched regions. These transcripts were more highly expressed in most mutants compared to wild type, especially if H4K20me3 is absent. **C)** RNA-seq reads of wild type and Δ*kmt5* mapped to accessory chromosome 17. The previous annotation (50) identified only a fraction of transcripts expressed on accessory chromosomes.

We noticed that, especially on accessory chromosomes, many regions without previously annotated genes or with incorrectly annotated genes showed transcriptional activity, often in a strand-specific manner and generating spliced transcripts, suggesting typical fungal transcription to generate mRNA by RNAPII. Many of these transcripts were more highly or newly expressed in the Δ*kmt5* mutant. To quantify the number of new transcripts that were not captured by the current gene annotation, we used stringtie (49) to identify transcripts in the combined RNA-seq data of all strains. We identified a total of 1,023 transcripts in facultative heterochromatin, of which only 516 were partially or fully overlapping with the previous annotation. Expression levels of these transcripts were highest in Δ*kmt5* and *hH4^K20M^* followed by Δ*ash1* (Figure 7B). The Δ*eaf3* and *hH4^K20/K20M^* strains exhibited only slightly higher expression of these transcripts (Figure 7B). Many of the newly discovered transcripts do not appear to encode for proteins, as they contain numerous, presumably premature, stop codons in all three reading frames. To better understand the potential function of these transcripts, we manually annotated all transcripts expressed on one accessory chromosome (chromosome 17), their chromatin state (in wild type), presence of additional copies of homologs on other accessory or core chromosomes, any potential function based on blastx searches, and presence of homologs in other species (Tables 2 and S2). We identified a total of 186 transcripts of which 55 match or partially match the previous annotation. Only 13 of these 186 transcripts seem likely to encode a protein based on the presence of an open reading frame without premature stop codons; one of these transcripts encodes a putative protease, while the others showed no predicted functional domains. If these 186 transcripts do not encode proteins, they may function as regulatory RNAs. We looked for homologous genes or sequences on other accessory chromosomes or core chromosomes using blastn searches in the *Z. tritici* wild-type genome. Almost half of the transcripts (89/186) on chromosome 17 had copies on other accessory chromosomes, often multiple copies on multiple chromosomes. Almost one third (62/186) showed similarity to sequences or genes on core chromosomes. Among the homologs on core chromosomes and functional annotation based on blastx searches, we found a striking enrichment of genes that encode for proteins involved in chromosome organization and segregation and cell cycle control. For example, we found eleven transcripts with similarity to the gene encoding a Shugoshin homolog (chr12_379), a protein that is important for chromosome cohesion during mitosis and meiosis. Seven transcripts are similar to those for protein kinases that are important for cell cycle control, one transcript has homology to transcripts of the kinetochore subunit, CenpC, two transcripts are similar to those of alpha tubulin, two transcripts are similar to those of two different Snf2 chromatin remodeling ATPases, and one transcript is similar to Sir2 histone deacetylase transcripts (Table S2). Half of these conserved transcripts are expressed as antisense mRNAs to the transcripts produced from their core chromosome homolog, while the other half is expressed from the same strand. Homologous genes on core chromosomes were not differentially expressed in any of the mutants, except for four genes (three in Δ*eaf3* and one in *hH4^K20M^*), but these do not seem to correlate with corresponding transcript expression levels on chromosome 17.

**Table 2:** Manual annotation of transcripts expressed on chromosome 17 based on RNA-seq data of wild type and mutants.

|  | number of transcripts |
| --- | --- |
| identified | 186 |
| protein coding | 13 |
| homologs on accessory chromosomes | 89 |
| homologs on core chromosomes | 62 |
| homologs in other (fungal) species | 41 |
| functional annotation | 31 |

## Discussion

We report chromatin dynamics and interactions in facultative heterochromatin that identify Kmt5 and H4K20me3 as important regulators for transcriptional repression and recruitment of Ash1-mediated H3K36me3, which itself at least partially regulates the distribution of H3K27me3. Here we investigated potential epistasis rules for repressive histone modifications and show that presence of H4K20me3 and Kmt5 are essential for Ash1-mediated H3K36me3, because we observed absence of H3K36me3 in selected regions upon either deletion of Kmt5 or introduction of a mutant histone H4^K20M^. We determined that H3K27me3 is controlled by both H4K20me3 and H3K36me3 levels, at least in most regions with H3K27me3 enrichment. Previously, no genetic or physical interactions between H3K36me3 and H4K20me3, or the methyltransferases involved (Ash1 and Kmt5, respectively) had been described. Our findings suggest that H4K20 methylation needs to be established before Ash1 can catalyze H3K36me3.

Based on our ChIP-seq, H4K20me3 is a widespread mark, heavily enriched along *Z. tritici* chromosomes, especially in regions that are enriched with H3K27me3, segments of the genome we refer to as facultative heterochromatin. Here we elucidated the function of H4K20me3 in the formation and maintenance of facultative heterochromatin. Why H4K20me3 is also enriched across some gene bodies outside of facultative heterochromatin remains unclear. In fungi, Kmt5 catalyzes all three methylation states, which makes it difficult to disentangle the relative importance of the three different potential H4K20 methylation states. By ChIP, we showed that H4K20me1 is not specifically enriched along the genome, but we showed that levels are lower in facultative heterochromatin, where H4K20me3 enrichment is highest, suggesting that these two methylation states do not coexist in facultative heterochromatin under the tested conditions. This makes the trimethyl state the most likely to be responsible for the effects we observed in facultative heterochromatin. We currently have no data on the distribution of H4K20me2.

In animals, H4K20 methylation is found in various chromatin environments, and in a tissue- and life stage-dependent manner (11). Because of the presence of multiple H4K20 methyltransferases, non-histone targets, and interdependency of methylation states, distinct roles for the different methylation states are difficult to study. H4K20me1 has been associated with both active (24, 51), as well as repressed genes (52), promoting chromatin openness (24), but is also enriched in inactive regions, such as the inactivated X-chromosome (53). Previous studies found that H4K20me3 levels increase during senescence and aged tissues but decrease in cancer cells, which is associated with both increased and decreased silencing, respectively, suggesting that H4K20me3 is important in suppressing tumorigenicity (54, 55). The C-terminus of KMT5C has been shown to interact with HP1, targeting H4K20me3 to H3K9me3-enriched regions, such as TEs, telomeres and pericentric regions (56, 57). This overlap of H4K20me3 with H3K9me3 is completely absent from *Z. tritici*, suggesting that Kmt5 has different interaction partners than in animals: H4K20me3 is not involved in silencing of constitutive heterochromatin in this species. In contrast to animals, fungi only possess a single methyltransferase mediating all H4K20me, as opposed to different enzymes catalyzing either H4K20me1 or H4K20me2/3. The presence of different H4K20 HMTs may result in more specialized interactions with different complexes that are specific to the different states of H4K20 methylation.

We found that H4K20me3 overlaps with Ash1-mediated H3K36me3 and H3K27me3 in facultative heterochromatin but also co-occurs with H3K36me3 outside of these regions. An overlap of H4K20me3 and H3K36me3 in gene bodies of actively transcribed genes has been shown in embryonic stem cells, possibly resembling the mechanisms that are responsible for H4K20me3 enrichment outside of facultative heterochromatin regions we found here (58). Very little is currently known about the connection between H4K20me and H3K27me3. H4K20me3 and H3K27me3 have been shown to co-occur on some transposons in *Xenopus tropicalis* embryos (59) and a similar recruitment mechanisms to X-chromosomes has been proposed for H4K20me1 and H3K27me3 (53). Results from other fungi are sparse, as no genome-wide studies have been carried out, even in model organisms such as *S. pombe* and *N. crassa*. Lastly, H4K20 methylation appears to be lacking completely in plants (60), suggesting that the control of H3K27me3-mediated facultative heterochromatin is regulated in entirely different ways.

To understand the biochemistry of a putative Kmt5 complex, we tagged the protein with GFP and short tags for protein purification. We showed that both N- and C-terminal tags greatly interfered with normal Kmt5 function, as documented by ChIP-seq. Taken together with our results on the added histone H4^K20M^ allele (H4^K20/K20M^), facultative heterochromatin may be the first region in which H4K20 methylation occurs, and from which this histone mark spreads into the bulk of the chromatin to normal levels observed in wild type. It is unclear how this partial activity occurs, but studies on other histone mutants, specifically H3^K27M^ showed that K to M mutations alter the release of histone methyltransferases from their substrates and thus sequester them on chromatin (61, 62). The consequence is limited or greatly reduced activity of the methyltransferase in other parts of the genome, which is what we observed with the H4^K20/K20M^ mutant. The reason N- and C-terminal Kmt5 fusion proteins show similar distribution may well relate to the overall structure of the Kmt5 protein, with the N-terminus being slightly more sensitive to modifications or additions than the C-terminus. In the absence of fully, or at least largely, complementing tagged alleles of Kmt5 we will address Kmt5 biochemistry in a separate study.

How then does H4K20 trimethylation govern facultative heterochromatin *de novo* assembly and maintenance? Our current working model (Figure 8) proposes that in maintenance mode either Kmt5 alone, or in a complex, methylates H4K20 in regions that are already enriched with methylated H4K20. For *de novo* methylation, some “signal” must be present. When Kmt5 is reintroduced into a Δ*kmt5* mutant, the protein, alone or in a complex, is able to detect such signals and methylate H4K20, as observed in the tagged Kmt5 strains we generated. These signals do not appear to be other histone modifications we analyzed in this study (H3K27me3, Ash1-mediated H3K36me3), as none of the mutants, except for Δ*kmt5* and histone *H4* mutants, reduced H4K20me3 levels in facultative heterochromatin. Analyses of putative DNA-based sequence motifs in regions enriched with H4K20me3 in the H4^K20/K20M^ mutant did not yield candidates.

**Figure 8:**
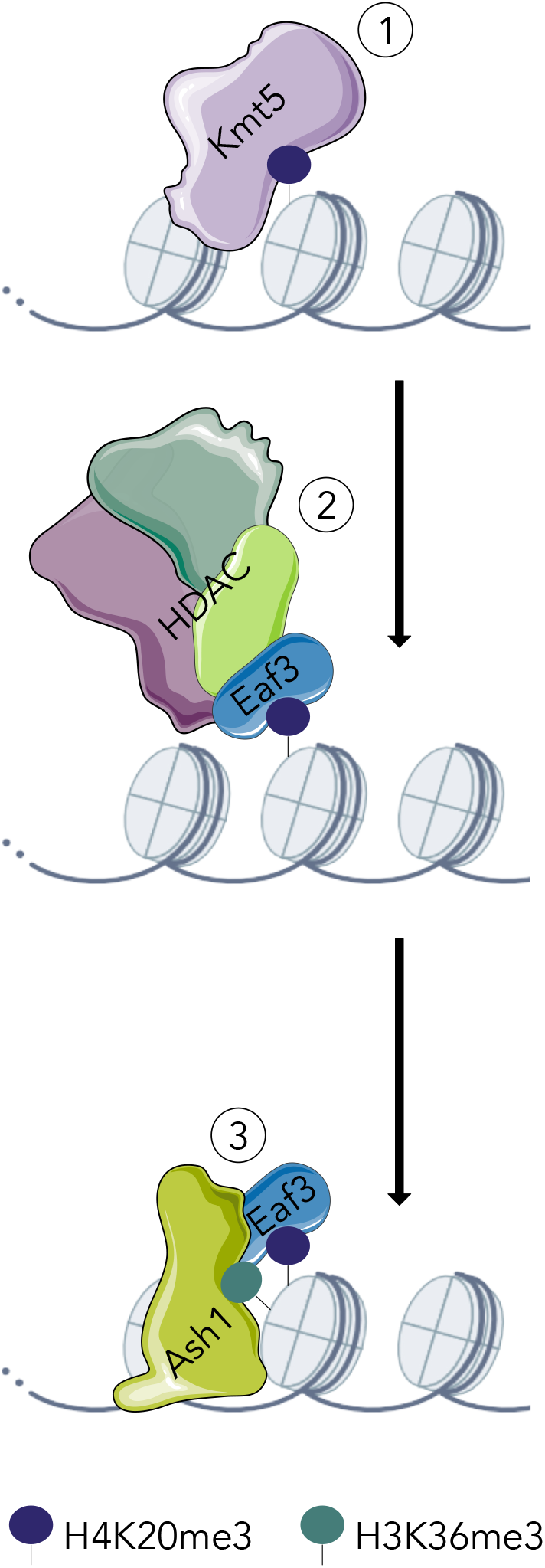
Model of how H4K20me3 is required for Ash1-mediated H3K36me3. **(1)** Kmt5 mediates H4K20me3 followed by **(2)** binding of Eaf3 to H4K20me3; Eaf3 may be in a complex with a histonedeacetylase (e.g., RPD3S). **(3)** Presence of Eaf3 promotes Ash1-mediated H3K36me3.

Independently from how Kmt5 first finds its targets, it seems likely that H4K20me3 is bound by Eaf3 which then stimulates activity of Ash1, as shown for MRG15 in flies (47, 63); this is supported by finding increased H4K20me3 levels in the absence of Eaf3 and the requirement of Eaf3 for Ash1-mediated H3K36me3. Whether Eaf3 is part of an Rpd3S-like complex in filamentous fungi, or whether it may directly recruit Ash1 to regions of H4K20me3 remains unknown. We propose that Ash-1-mediated H3K36 methylation results in a switch of Eaf3 binding from H4K20me3 to H3K36me3, though it is also possible that it may bind both modifications at the same time.

Little is known about proteins that interact with Kmt5 in fungi except for *S. pombe* Pdp1 (42). Here we showed that there is no true ortholog for this protein in filamentous fungi, and that the best homolog in *Z. tritici* and other filamentous fungi is an ortholog of PWWP domain *S. cerevisiae* Ioc4 protein, a component of some ISWI chromatin remodeling complexes in *N. crassa* (43). While this protein does not seem to be important for H4K20me3, we did observe increased enrichment of H3K27me3 outside of wild-type facultative heterochromatin regions, suggesting an important role in chromatin regulation; modest increases of H3K27me3 enrichment was also observed in *N. crassa* (43).

Similar to findings in *N. crassa* and *F. fujikuroi* but unlike in animals, H3K27me3 and Ash1-mediated H3K36me3 occur in the same regions (28, 30, 31, 36). Loss of Ash1-mediated H3K36me3 has minor impacts on H3K27me3 in *Z. tritici*, resulting in increased enrichment in some regions and decrease in other regions. While Ash1 in animals is predominantly a H3K36me2 methyltransferase, we found major impacts on H3K36me3, as shown previously in *N. crassa* and *F. fujikuroi* (30, 31). The fungal Ash1 proteins are different from Ash1 in animals, as they lack the C-terminal Bromo, PHD and BAH domains, suggesting they have different chromatin binding properties and thus functions than their animal homologs.

Functional analyses of the Kmt5 homolog in *S. pombe*, Set9, and some of the animal Kmt5 proteins suggest importance during replication and DNA repair (15, 16, 20, 21, 64, 65). Here we showed that both Kmt5 and Ash1 are necessary, but not essential, for normal growth of *Z. tritici*. While the Δ*ash1* mutant showed only minor effects under our conditions, the Δ*kmt5* mutant was more sensitive to genotoxic stress, especially at higher temperatures. Lack of both Kmt5 and Ash1 also resulted in accelerated chromosome loss rates: While the loss rate for wild type was ~3%, mirroring our earlier results (32, 37), loss of accessory chromosomes was 41% and 58% in Δ*ash1* and Δ*kmt5*, respectively. Increased chromosome loss affected some chromosomes more than others. Chromosomes 14 and 16 are easily lost even in wild-type strains, especially under heat stress conditions (32, 35, 37), and they were also lost most commonly, by far, in Δkmt5 and Δash1. Thus, unlike the loss of *kmt6*, which results in an overall more stable genome (32), lack of *ash1* and *kmt5* both resulted in decreased genome stability, which suggests different, separable functions in genome maintenance and control of gene expression in facultative heterochromatin for both Ash1 and Kmt5.

Our earlier studies showed that lack of H3K27me3 alone did not result in large-scale upregulation of regions enriched with H3K27me3 (32). One of our goals was to uncover which *Z. tritici* histone modifications, if any, take on the gene silencing function usually mediated by H3K27me3 in other organisms. Here, we showed that H4K20me3, and Ash1-mediated H3K36me3 are important for transcriptional repression of facultative heterochromatin. Some of these effects are phenocopied by replacement of histone H4K20 with a methionine (H4^K20M^), or in a strain with a wild type and H4K20M allele (H4^K20/K20M^). While some *bona fide* genes on core chromosomes and even accessory chromosomes are affected by the lack of Kmt5 or Ash1, most regions enriched with H3K27me3, Ash1-mediated H3K36me3, and H4K20me3 have relatively few previously annotated genes. To understand the transcripts that emanate from these regions we manually curated RNA-seq data for one accessory chromosome. We found that most transcripts arise from sequences that do not appear to be regular genes. How exactly these RNA species act, what their specific functions may be, and whether they interfere or aid propagation of accessory chromosomes in *Z. tritici* goes beyond the scope of the current study. Our results show, however, that studies resulting in a lack of specific histone modifications may necessitate renewed annotation efforts, which should capture transcripts from pseudogenes and apparent *bona fide* genes with few nonsense codons.

In conclusion, we showed that H4K20me3, H3K27me3, and Ash1-mediated H3K36me3 delineate facultative heterochromatin in *Z. tritici*. H4K20me3, as well as the presence of Eaf3, are essential for Ash1-mediated H3K36me3, uncovering a novel mechanism for establishment and maintenance of facultative heterochromatin and transcriptional silencing.

## Methods

### Fungal and bacterial growth conditions

*Zymoseptoria tritici* cultures were grown on YMS (4 g yeast extract, 4 g malt, 4 g sucrose per liter, with 16 g agar per liter added for solid medium). Glycerol stocks (1:1 YMS and 50 % glycerol) were maintained at −80°C. *Escherichia coli* strain DH10beta (NEB) was grown at 37°C in LB (10 g tryptone, 10 g NaCl, 5 g yeast extract per liter) with appropriate antibiotics (100 μg/mL kanamycin). *Agrobacterium tumefaciens* strain AGL1 was grown in dYT (16 g tryptone, 5 g NaCl, 10 g yeast extract per liter) or LB media with appropriate antibiotics (100 μg/mL kanamycin, 100 μg/mL carbenicillin, 50 μg/mL rifampicin).

### Generation of plasmids for fungal transformation

Plasmids were generated by Gibson Assembly (66) using a homemade Gibson reaction mix based on recipes from OpenWetWare (OpenWetWare, https://openwetware.org/wiki/Gibson_Assembly) (5x Isothermal Reaction Mix [600 μL]: 300 μL 1 M Tris-HCl [pH 7.5], 30 μL 1 M MgCl2, 6 μL 100 mM dGTP, 6 μL 100 mM dATP, 6 μL 100 mM dTTP, 6 μL 100 mM dCTP, 30 μL 1 M DTT, 0.15 g PEG 8000, 60 μL 50 mM NAD^+^, 156 μL H_2_O) and modified by using the “enhanced” version (68) (1.33x Gibson-Assembly Master-Mix [for 25 reactions]: 100 μL 5x Isothermal Reaction-Mix, 215.32 μL H2O, 0.2 μL T5 Exonuclease [10 U/μL NEB #M0363], 6.25 μL Phusion DNA-Polymerase [2 U/μL ThermoFisher #F530L], 3.125 μL ET SSB [500 μg/ml NEB #M2401S], 50 μL Taq DNA-Ligase [40 U/μL stock NEB #M0208L]). Primers, plasmids, and strains used in this study are listed in Table S3.

### Transformation of *Zymoseptoria tritici*

Transformation was carried out by *A. tumefaciens* (69). In our updated protocol (detailed protocol is provided as supplemental method), *A. tumefaciens* strain AGL1 (transformed with the respective target plasmid, see Table S3) was grown for ~24 h in liquid culture (LB with 100 μg/mL kanamycin, 100 μg/mL carbenicillin, and 50 μg/mL rifampicin) at 28°C before induction. Induction was performed by centrifuging LB cultures (3000rpm for 5 min), and resuspending cells in 1 mL of induction medium (10 mM K_2_HPO_4_, 10 mM KH_2_PO_4_, 2.5 mM NaCl, 2 mM MgSO_4_, 9 μM FeSO_4_, 40 mM MES buffer, 0.7 mM CaCl_2_, 0.5 % glycerol, 5 mM glucose, 6.25 mM NH_4_NO_3_, 0.002 % Vogel trace elements, 100 μg/mL kanamycin, 100 μg/mL carbenicillin, 50 μg/mL rifampicin, 200 μM acetosyringone) and inoculating resuspended AGL1 cultures in 5 mL induction medium (OD_590_ = 0.15) for overnight growth at 28°C. Induced cultures were adjusted to OD_590_ = 0.4 the following day and mixed with *Z. tritici* cells (OD_590_ = 1) at a 1:1 ratio. To grow recipient *Z. tritici* cells for this step, spores were streaked out on YMS plates from glycerol stocks 3-4 days before transformation, resuspended in water, and the concentration adjusted to an OD_590_ = 1 in induction medium, before mixing with induced AGL1 cells. Aliquots of 250 μL of the *A. tumefaciens* AGL1 and *Z. tritici* mixture were plated on Hybond N+ membranes on plates with induction medium and incubated at room temperature for 48-72 h. Membranes were transferred to YMS plates containing the appropriate antibiotics for selection (250 μg/mL cefotaxime, 150 μg/mL timentin, 100 μg/mL G418, 100 μg/mL hygromycin) and incubated for another ~10 days at 18°C until resistant colonies appeared.

### Chromatin immunoprecipitation and sequencing (ChIP-seq)

ChIP experiments were carried out as described previously (70) with some modifications; a detailed protocol is provided as supplemental method. Briefly, fungal spores were grown on YMS plates for three to four days, harvested, and resuspended in 5 mL of 1× PBS (137 mM NaCl, 2.7 mM KCl, 10 mM Na_2_HPO_4_, 1.8 mM KH_2_PO_4_). Cells were crosslinked using 0.5% formaldehyde for 15 min at room temperature and quenched by adding 150 μL of 2.5 M glycine. Crosslinked cells were centrifuged at 3,000 rpm for 5 min at 4°C and washed once with cold 1× PBS. We found that grinding cell pellets to a fine powder in liquid nitrogen and storage at −80°C resulted in no noticeable decrease in ChIP efficiency (storage for at least six months was acceptable). For ChIP, frozen, ground cells were resuspended in ice-cold ChIP lysis buffer (50 mM HEPES-NaOH [pH 7.5], 90 mM NaCl, 1 mM Na-EDTA [pH 8.0], 1% Triton X-100, 0.1% DOC, and proteinase inhibitors [1 mM PMSF, 1 μg/mL Leupeptin, 1 μg/mL E-64 and 0.1 μg/mL Pepstatin]) in a ratio of 5 μL ChIP lysis buffer to 1 mg of ground cells. To 1 mL of lysate, 2 μL 1 M CaCl_2_ and 5 μL micrococcal nuclease (NEB, #M0247S) were added and chromatin was fragmented into predominantly mono-nucleosomes by incubation at 37°C for 15 min. The reaction was stopped by adding 20 μL of 0.5 M Na-EGTA, pH 8. Digested chromatin was centrifuged at 6,000 rpm for 5 min and 250-300 μL of the supernatant was used per ChIP reaction. Chromatin was pre-cleared by incubating with protein A Dynabeads (Invitrogen) at 4°C for 1 h followed by overnight incubation with 3 μL of the following antibodies: H3K4me2 (Millipore 07-030), H3K9me3 (Active Motif 39161), H3K27me3 (Active Motif 39155), H3K27me2 (Active Motif 39245), H3K36me2 (Active Motif 39255; abcam 9049), H3K36me3 (Active Motif 61101; abcam 9050), H4K20me1 (abcam 9051), H4K20me3 (Active Motif 91107). After overnight incubation, 25 μL protein A Dynabeads were added and samples were incubated for 1-2 h at 4°C. Beads were washed twice with ChIP lysis buffer, once with ChIP lysis buffer + 0.5 M NaCl (50 mM HEPES-NaOH [pH 7.5], 500 mM NaCl, 1 mM Na-EDTA [pH 8.0], 1% Triton X-100, 0.1% DOC), once with LiCl wash buffer (10 mM Tris-HCl [pH 8.0], 250 mM LiCl, 0.5 % IGEPAL CA-630, 0.5 % DOC, 1 mM Na-EDTA [pH 8]) and once with 1x TE buffer. Chromatin was eluted in 125 μL TES (50 mM Tris-HCl [pH 8.0], 10 mM Na-EDTA [pH 8], 1% SDS) and de-crosslinked overnight at 65°C. Samples were treated with RNAse A (2 h at 50°C) and proteinase K (2 h at 65°C) and DNA was extracted with the ChIP DNA Clean & Concentrator kit (Zymo Research). Libraries for sequencing were prepared with a modified version of the Next Ultra II DNA Library Prep Kit for Illumina (NEB, #E7645S). Sequencing was performed on an Illumina HiSeq 3000, obtaining 2 × 150-nt read pairs by Admera Health (South Plainfield, NJ, USA).

### RNA isolation and sequencing (RNA-seq)

Fungal cells were grown and harvested as described for the ChIP experiments. Cells were ground in liquid nitrogen and ~100 mg of ground cells were used for total RNA extraction by a TRIZOL (Invitrogen) method. Extraction was carried out according to the manufacturer’s instructions and total RNA was treated with DNAse I. Stranded mRNA libraries were prepared by a modified version of a previously published protocol (71); the detailed protocol is provided as supplemental method. Stranded mRNA libraries were sent to Admera Health (South Plainfield, NJ, USA) for sequencing on an Illumina HiSeq 3000, obtaining 2 × 150-nt read pairs.

### Quantification of chromosome loss

Each strain (wild type, Δ*kmt5*, Δ*ash1*) was grown in five replicate populations on YMS plates, and every three to four days a fraction of the population was transferred to a new plate. After eight transfers (four weeks), populations were diluted and spread on YMS plates to obtain single colonies, representing individual cells of the respective population. A total of 48 colonies/replicate (i.e., 240 per strain) were screened by PCR for the presence of all accessory chromosomes as previously described (37).

### Western blots

Fungal cells were grown on YMS plates for three to four days, harvested and washed in 1x PBS. Cells were resuspended in three volumes of high-salt extraction buffer (25 mM HEPES-NaOH [pH 7.9], 700 mM NaCl, 0.1 mM Na-EDTA [pH 8.0], 0.2% NP-40, 20% glycerol, 1.5 mM MgCl_2_, 1 mM PMSF, 1 mM DTT, 1 μg/mL Leupeptin, 0.1 μg/mL Pepstatin) and incubated on ice for 10 min. Samples were sonicated (Branson Sonifier-450) for three sets of 10 pulses (output = 2, duty cycle = 80), keeping the sample on ice between sets. Insoluble material was removed by centrifugation at 14,000 rpm and 4°C for 10 min. Protein concentrations were quantified with a Qubit fluorimeter (Invitrogen). Total protein (75 μg per lane) was loaded onto pre-cast 4-20% gradient Mini-PROTEAN TGX Stain-Free SDS PAGE gels (BioRad). Proteins were transferred onto nitrocellulose membrane (0.2 μm pore size; BioTrace) by wet transfer (1 h, 110V). The following primary antibodies were used: ⍺H3 (Active motif 39763, mAb, 1:5000), ⍺H4K20me1 (abcam 9051, pAb, 1:1000), ⍺H4K20me3 (Active motif 39180, pAb, 1:2000). The following secondary antibodies were used: IRDye 680RD Goat anti-Mouse IgG (LI-COR 926-68070, 1:5000), IRDye 800CW Goat anti-Rabbit IgG (LI-COR 926-32211, 1:5000). Fluorescent signals were detected using a LI-COR Odyssey CLx imager.

### Hi-C analyses

WT and Δ*kmt6* cells were grown as described for ChIP experiments. Hi-C experiments were carried out using the Phase Genomics Proximo Hi-C Kit (Microbe) following the instructions for fungal samples; the Proximo Hi-C Kit (Fungal) had not been officially released when we carried out our experiments. *Dpn*II was used as restriction enzyme. Libraries were generated and size-selected according to the Phase Genomics kit instructions. Libraries were sent for sequencing to Phase Genomics, obtaining ~100 million 2 × 100-nt reads per sample.

### Phenotypic responses to genotoxic stress

Fungal cells were grown on YMS plates for three to four days and diluted to OD_590_ = 1 in water (~10^7^ cells/mL and tenfold dilution series to 100 cells/mL); 3 μL of the spore dilutions were pipetted onto plates. Pictures were taken after 7 and 12 days of growth (18°C, 28°C, and RT [~23°C]). To test for responses to different genotoxic stressors *in vitro*, YMS plates containing hydroxyurea (HU; 100 μM, 1 mM, 10 mM), camptothecin (1 mM, 10 mM, 100 mM), and methyl methanesulfonate (MMS; 0.005% and 0.0075%) and three plates containing only YMS were prepared.

### Data analyses

Detailed descriptions of software and commands for analyses of ChIP-seq, RNA-seq and Hi-C data can be found as supplementary methods. A list of all sequencing datasets can be found in Table S4.

### ChIP-seq

ChIP-seq data were quality-filtered and adapters removed with trimmomatic v.0.39 (72). Mapping was performed with bowtie2 v.2.4.4 (73), and sorting and indexing with samtools v.1.9 (74). Normalized coverage bigwig files and heatmaps were created with deeptools v.3.5.1 (75). Wiggletools v.1.2 and the UCSC Genome Browser tools were used to calculate means for replicates and converting wig to bigwig files. Peaks were called with homer v4.11.1 (76) and bedtools v2.30.0 were used to make windows and intersect peaks with genomic features.

### RNA-seq

RNA-seq data were quality filtered and adapters removed with trimmomatic v.0.39 (72). Mapping of all paired reads was performed with hisat2 2.1.0 (77), and sorting and indexing with samtools v.1.9 (74). HTSeq v.0.11.2 (78) was used to count reads on features. The *Z. tritici* IPO323 gene annotation was obtained from FungiDB (release 53) (79), based on previous annotations (80). Differential expression analyses was conducted with DEseq2 v.1.32.0 (81), see supplemental R script. New transcripts were annotated by combining all mapped RNA-seq data with stringtie v. v2.2.1 (49). Manual annotation was done by combining all RNA-seq data and extracting transcript sequences that were > 200 bp in length and supported by at least 10 reads with IGV (82) and Geneious (83). Transcripts were characterized by blastn with the *Z. tritici* genome as a reference to find homologous sequences on core and accessory chromosomes and blastx to find homologs in other species and to infer functional annotations.

### Hi-C

HiCUP v0.8.3 (84) was used to trim and map the data, see supplemental config file. Homer v4.11.1 (76) was used to extract valid Hi-C interactions and generating normalized interaction matrices. Treeview3 (85) was used to visualize Hi-C data.

## Supporting information

Suppl_Method_IlluminaLib

Suppl_Methods_RNAseq

Suppl_Methods_HiC

Suppl_Methods_ChIP

Suppl_Method_Transformation

Suppl_Method_RNALib

Suppl_Method_ChIPZymo

## Data availability

Sequencing raw reads (FASTQ files) of all ChIP-seq, RNA-seq and Hi-C data are available online at Sequence Read Archive (SRA) under BioProject ID PRJNA902413.

Normalized bigwig files for all ChIP-seq datasets, reference genome and annotations have been deposited at zenodo under doi: 10.5281/zenodo.7460287.

## Acknowledgements

We thank Eva Stukenbrock for supplying critical materials and strains, and for supporting this work while MM was still at CAU Kiel. We thank colleagues in the Freitag lab and Zachary Lewis (University of Georgia) for conversations and comments on the manuscript. MM was supported by a grant from the DFG (MO 3755/1-1). JBR was supported by a grant from USDA-AFRI (2019-67012-29722). Chromatin research in the Freitag lab is supported by grants from the NSF (MCB1818006), NIH (R01GM132644), and the BSF (#2019034) to MF.

## Supplement

### Tables

**Table S1**. DESeq2 results for all mutants in comparison to wild type including functional annotation.

**Table S2**. Detailed overview of all manually annotated transcripts on chromosome 17.

**Table S3**. List of all plasmids, strains and primers generated and used in this study.

**Table S4**. Summary of all sequencing datasets generated in this study.

---

**Figure S1.**
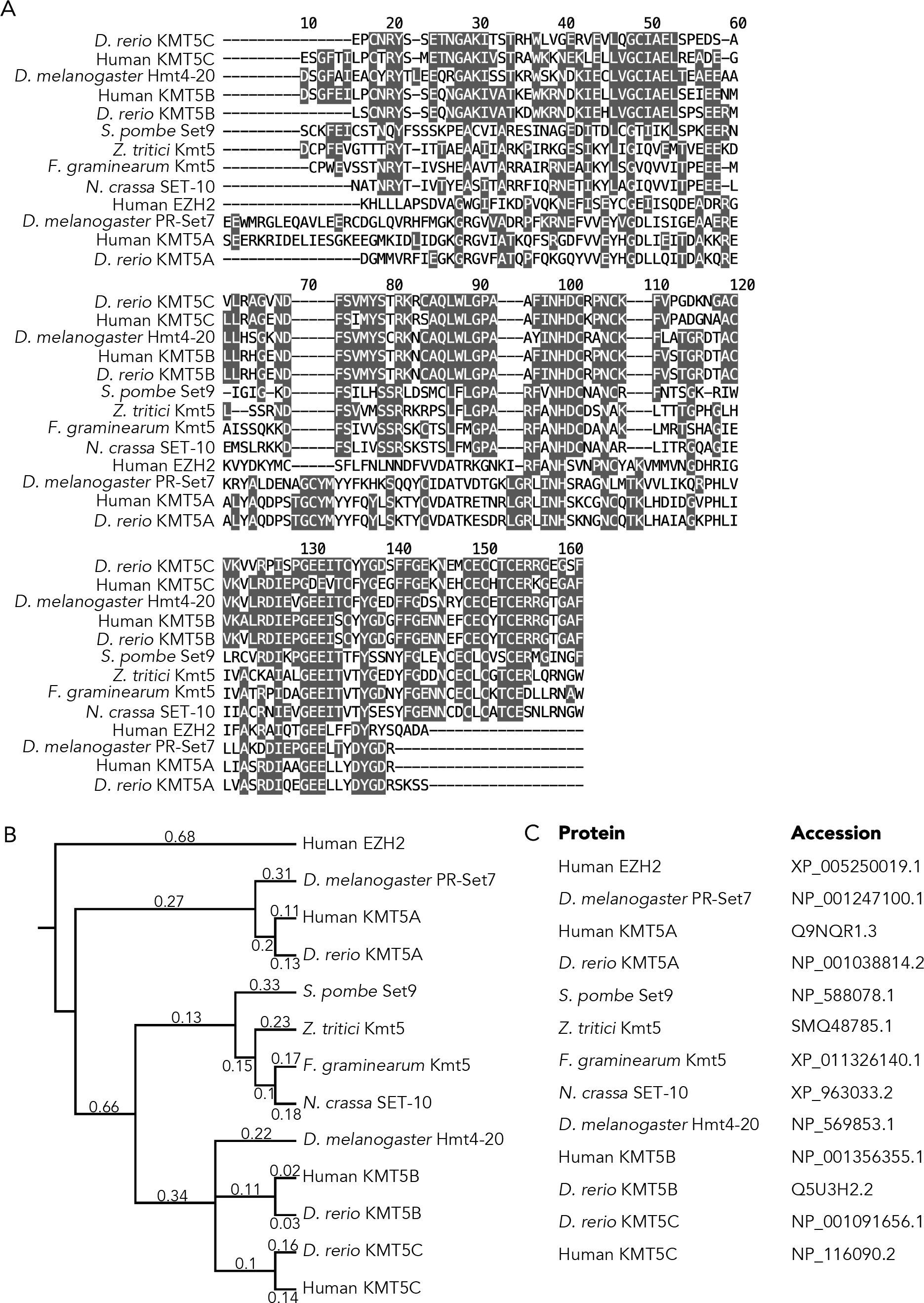
Comparison of H4K20 HMT SET domains. A) Alignment of SET domains of H4K20 methyltransferases in different species. SET domain of human EZH2 is used as an outgroup. B) Phylogenetic tree based on the alignment in A. Numbers indicate substitutions per site. C) Accession numbers of proteins.

**Figure S2.**
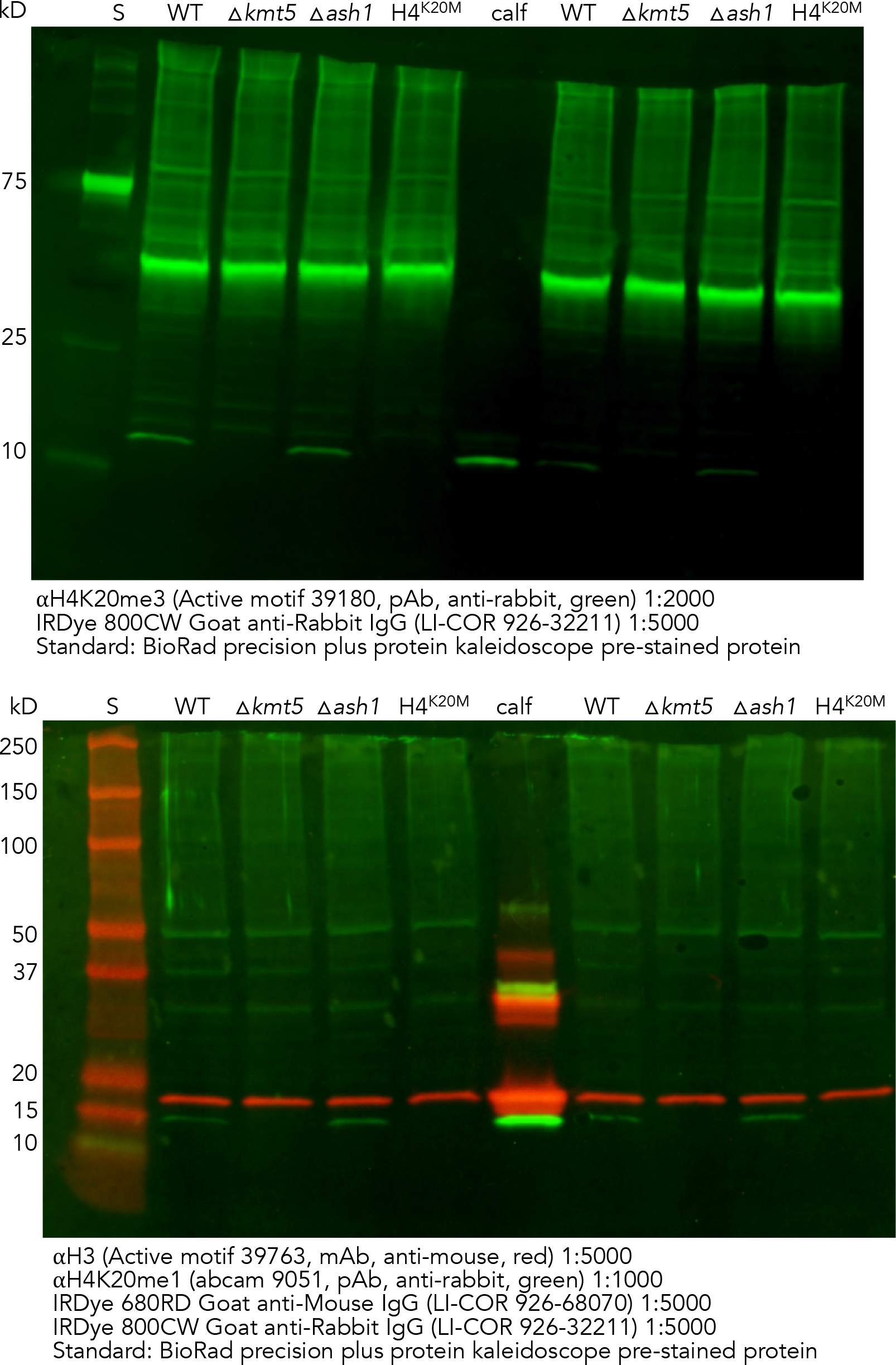
Uncropped western blot images for detection of H4K20me1, H4K20me3 and H3.

**Figure S3.**
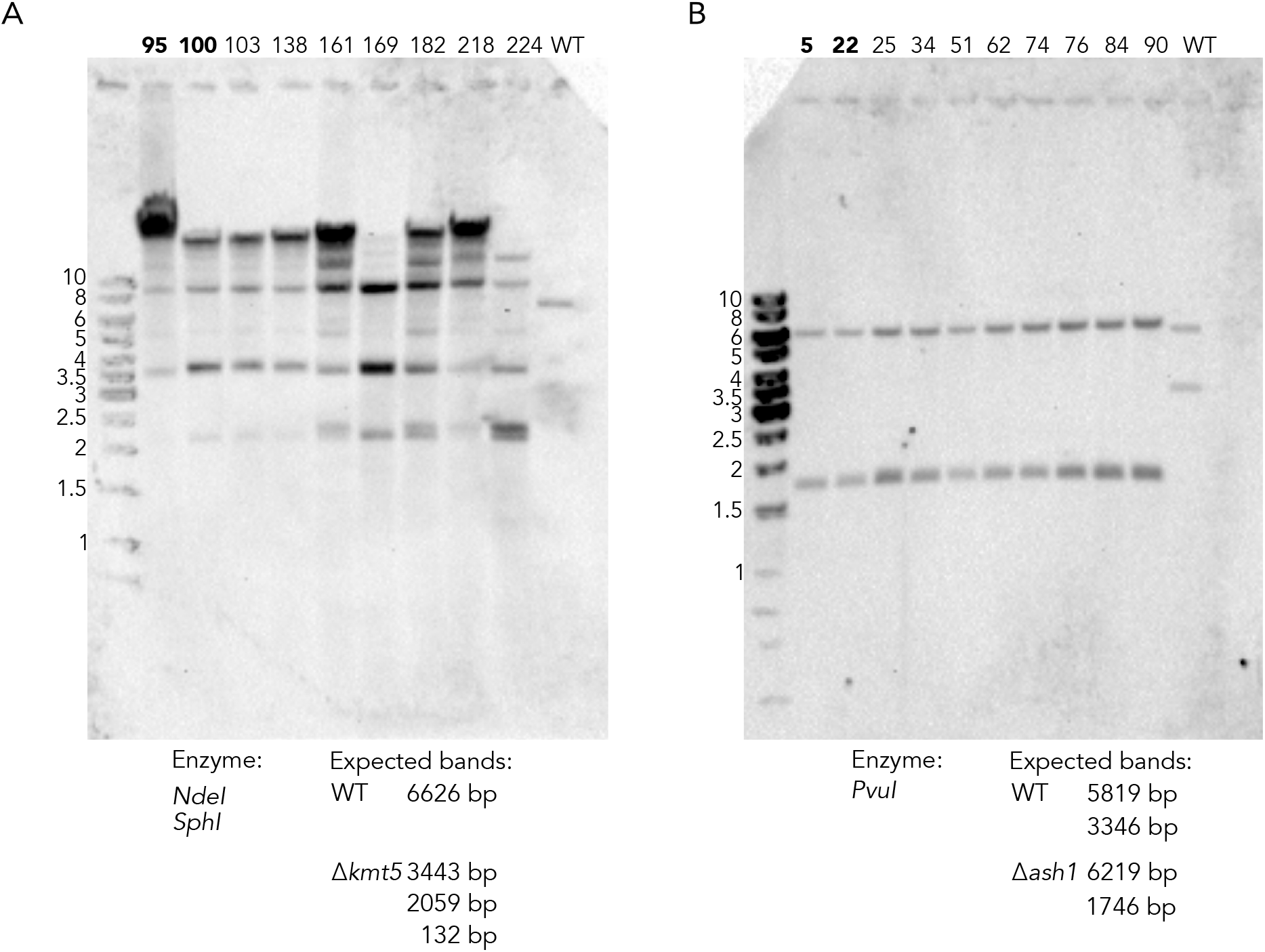
Southern blots to confirm deletion of *kmt5* and *ash1*. A) Δ*kmt5* mutants show more bands in addition to the expected bands. Because one of the flanks used for homologous recombination and as probe for Southern analyses is a repetitive element, integration into the repetitive element may have altered restriction sites, and additional bands are detected because of TE copy number. Deletion of *kmt5* was confirmed in all ChIP- and RNA-seq datasets. Strains used for further analyses were 95 and 100. B) Confirmation of Δ*ash1*. Strains used for further analyses were 5 and 25.

**Figure S4.**
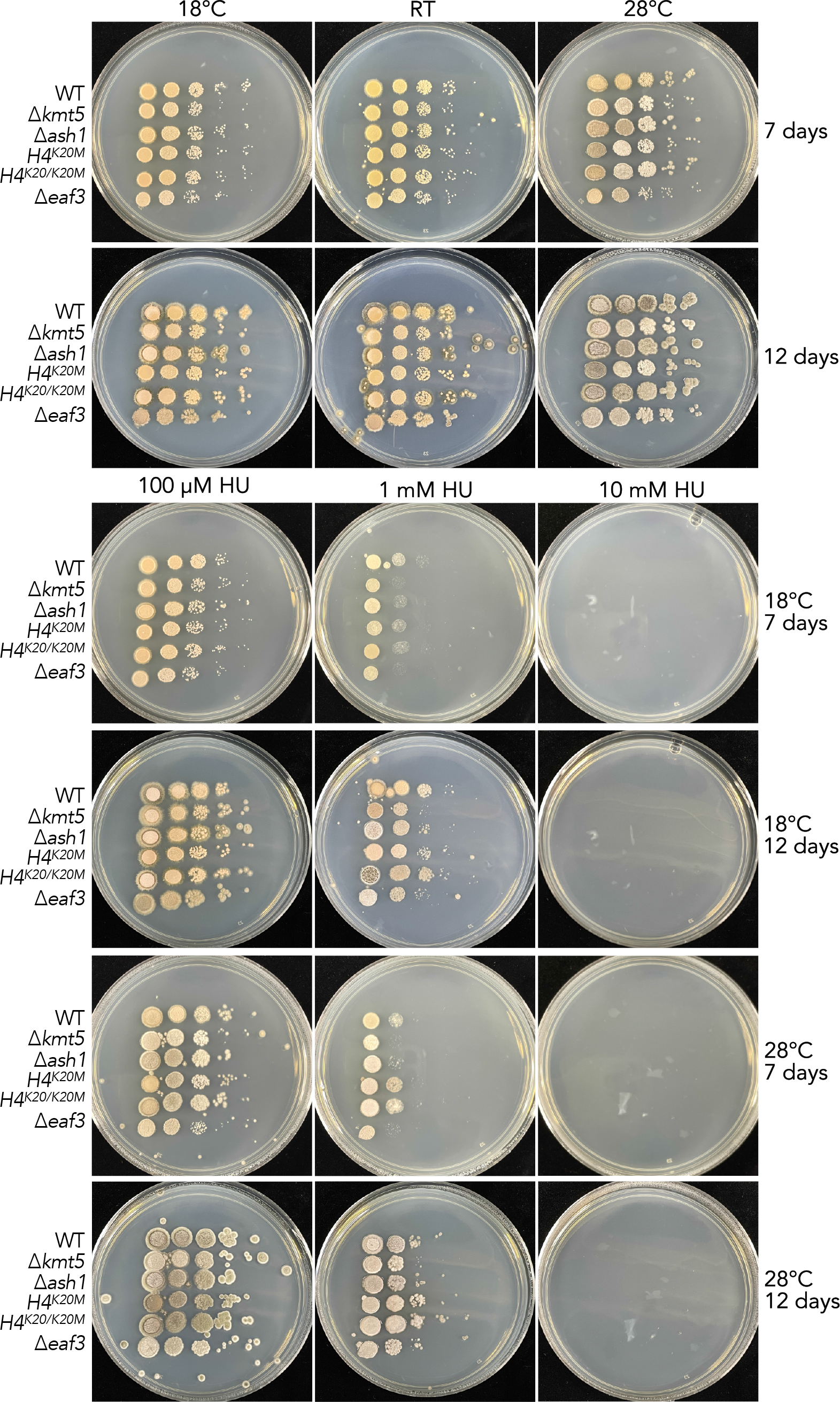
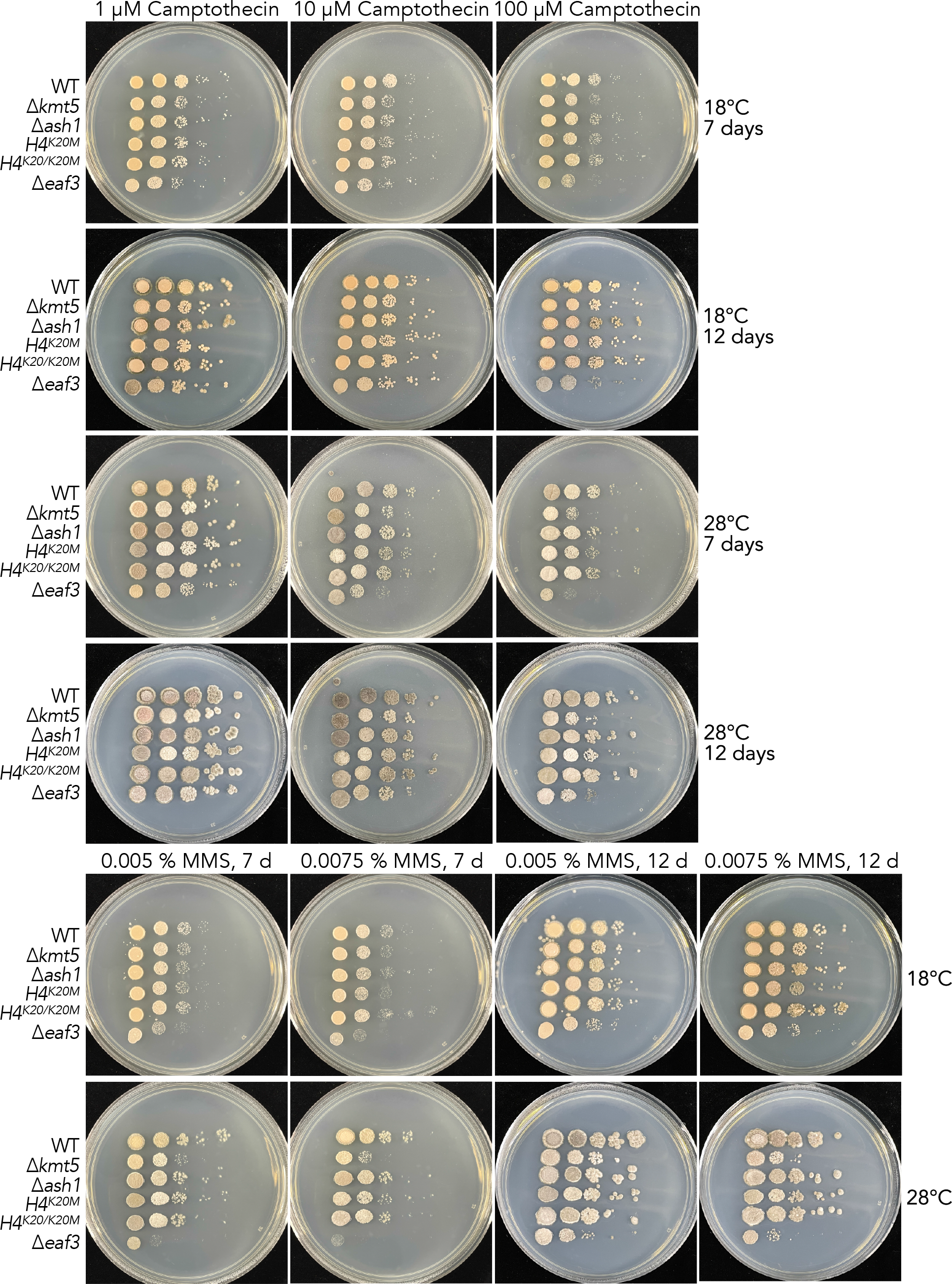
Phenotypic characterization of WT, Δ*kmt5*, Δ*ash1*, *H4^K20M^*, H4^K20/K20M^ and Δ*eaf3* under different genotoxic stress conditions and temperatures.

**Figure S5.**
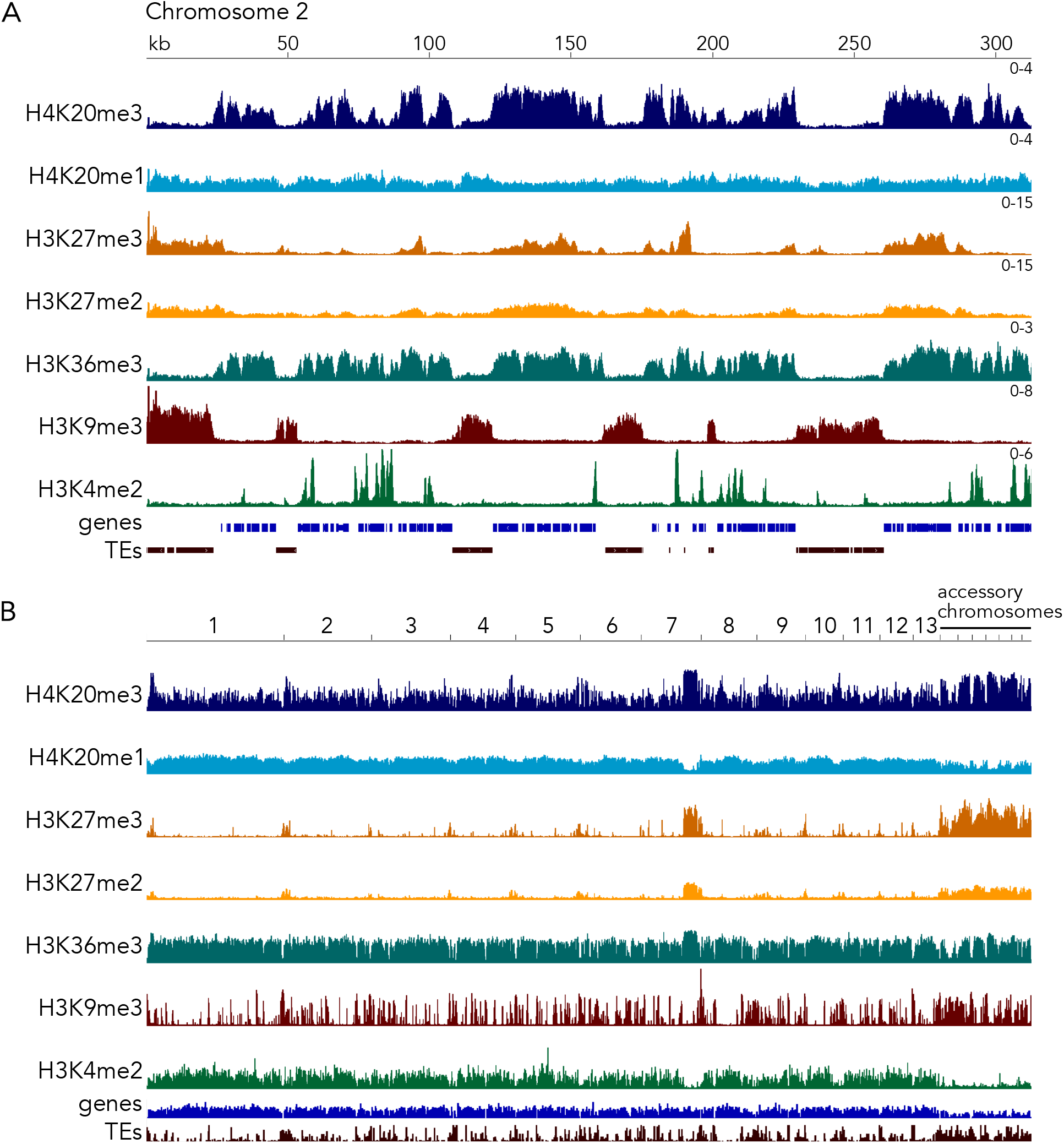
Genome-wide distribution of H4K20me3, H3K36me3, H3K27me3, H3K9me3 and H3K4me2 in wild type.

**Figure S6.**
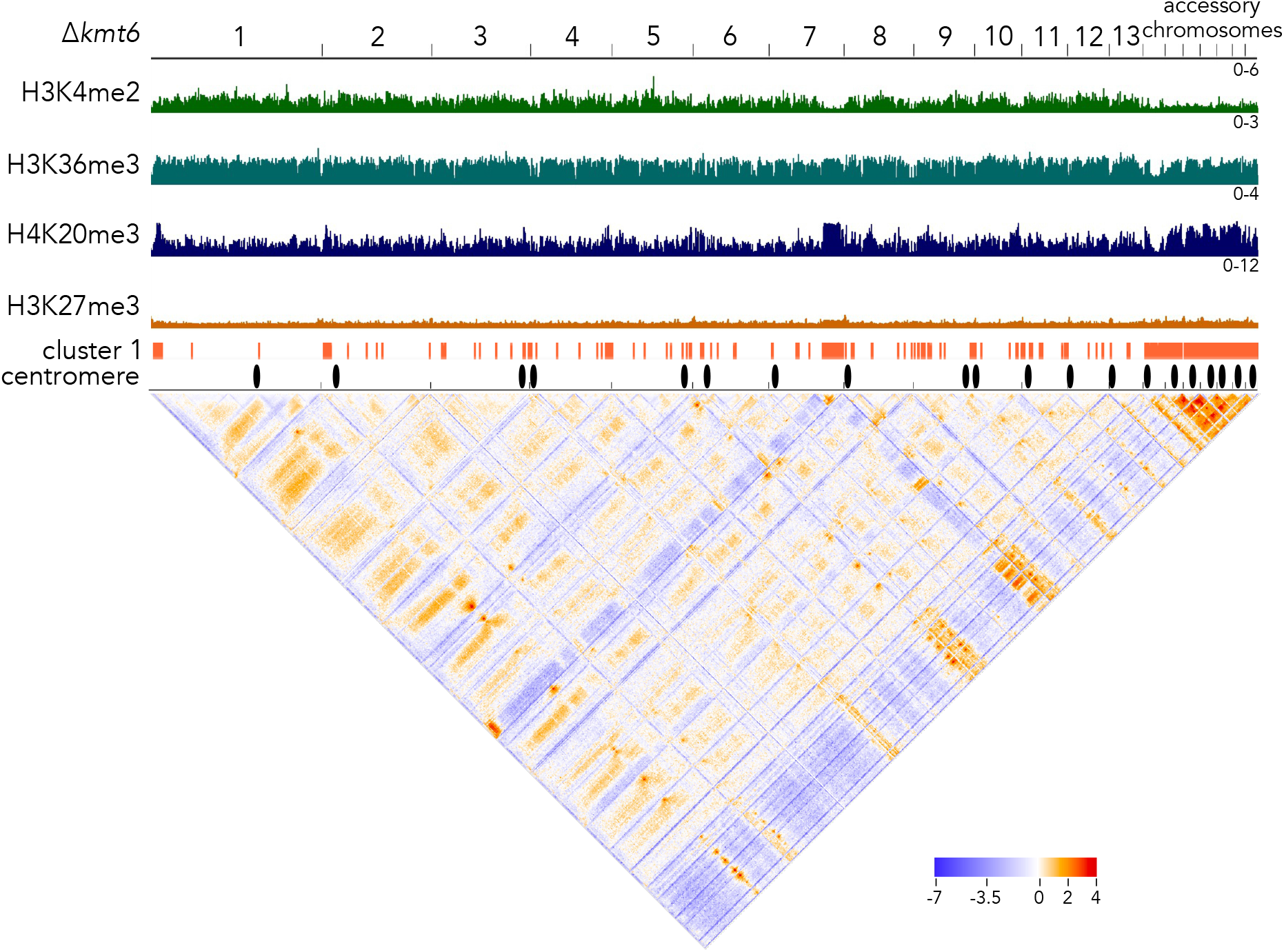
Interacting genomic regions in the Δ*kmt6* genome. Shown are log_2_ ratio of observed interactions relative to expected interactions, fewer interactions than expected are shown in blue, more interactions are shown in red.

**Figure S7.**
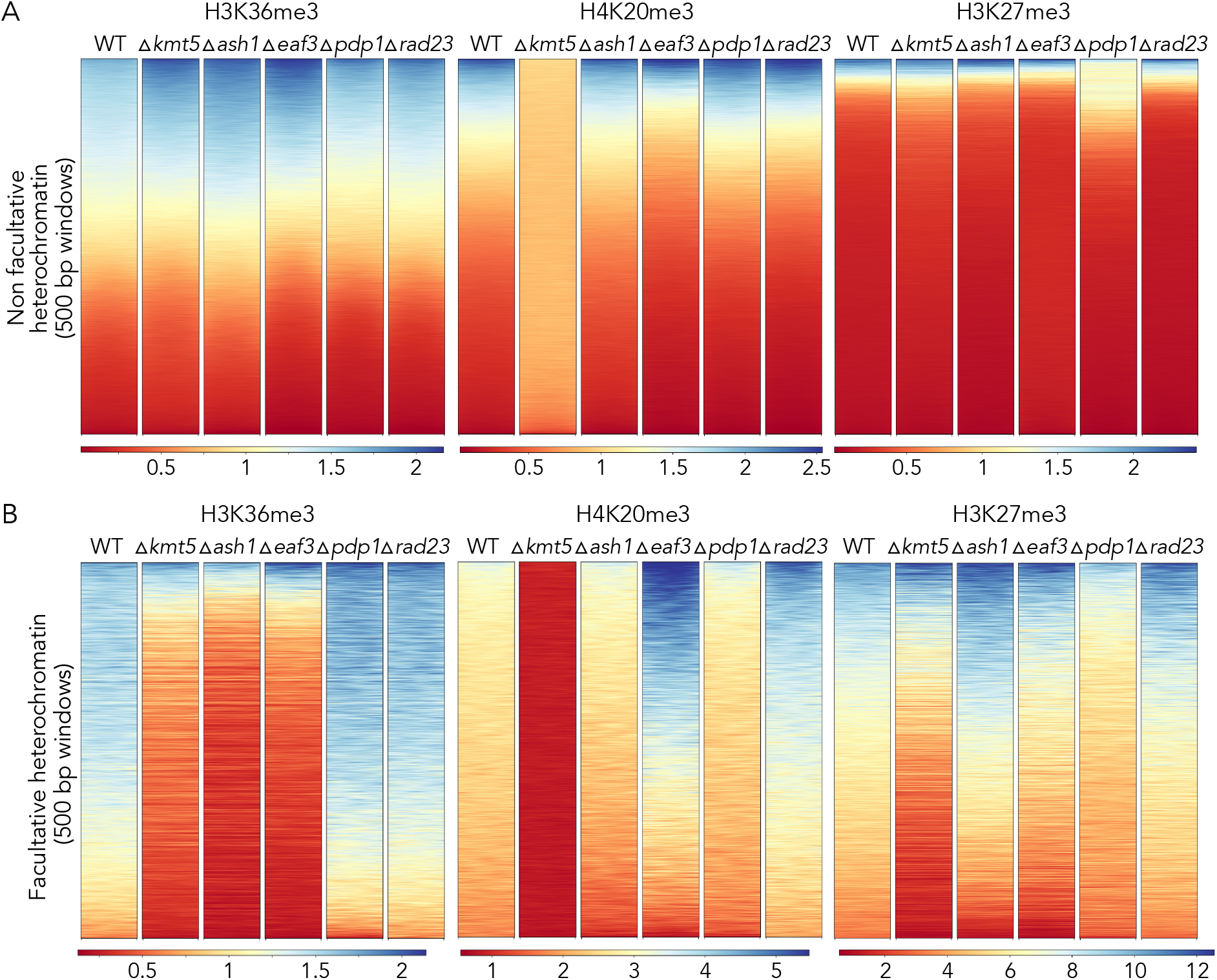
Enrichment of different histone marks outside (A) and within (B) facultative heterochromatin regions. While we observed subtle changes outside of facultative heterochromatin, we observed the most prevalent differences in regions of facultative heterochromatin.

**Figure S8.**
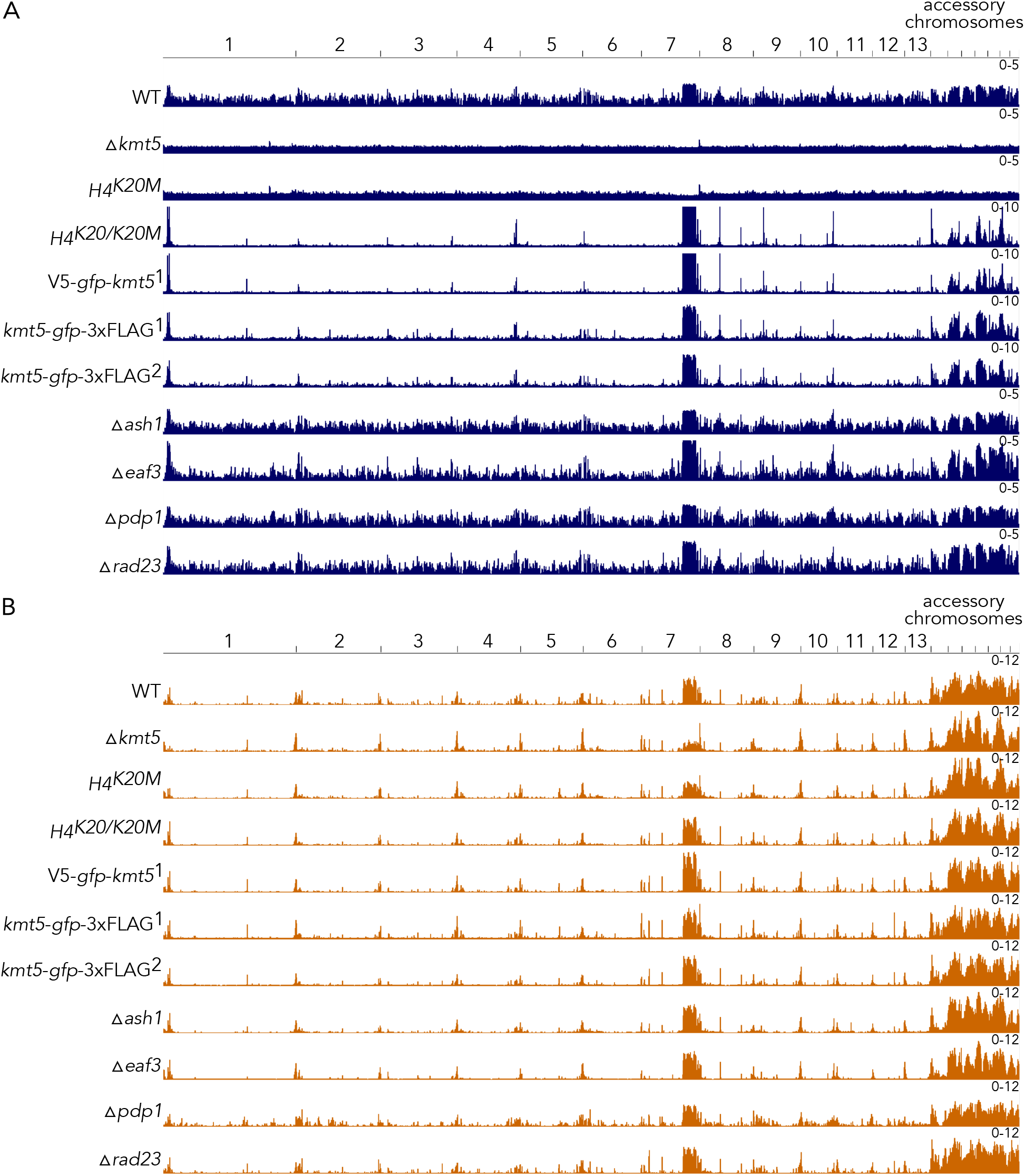
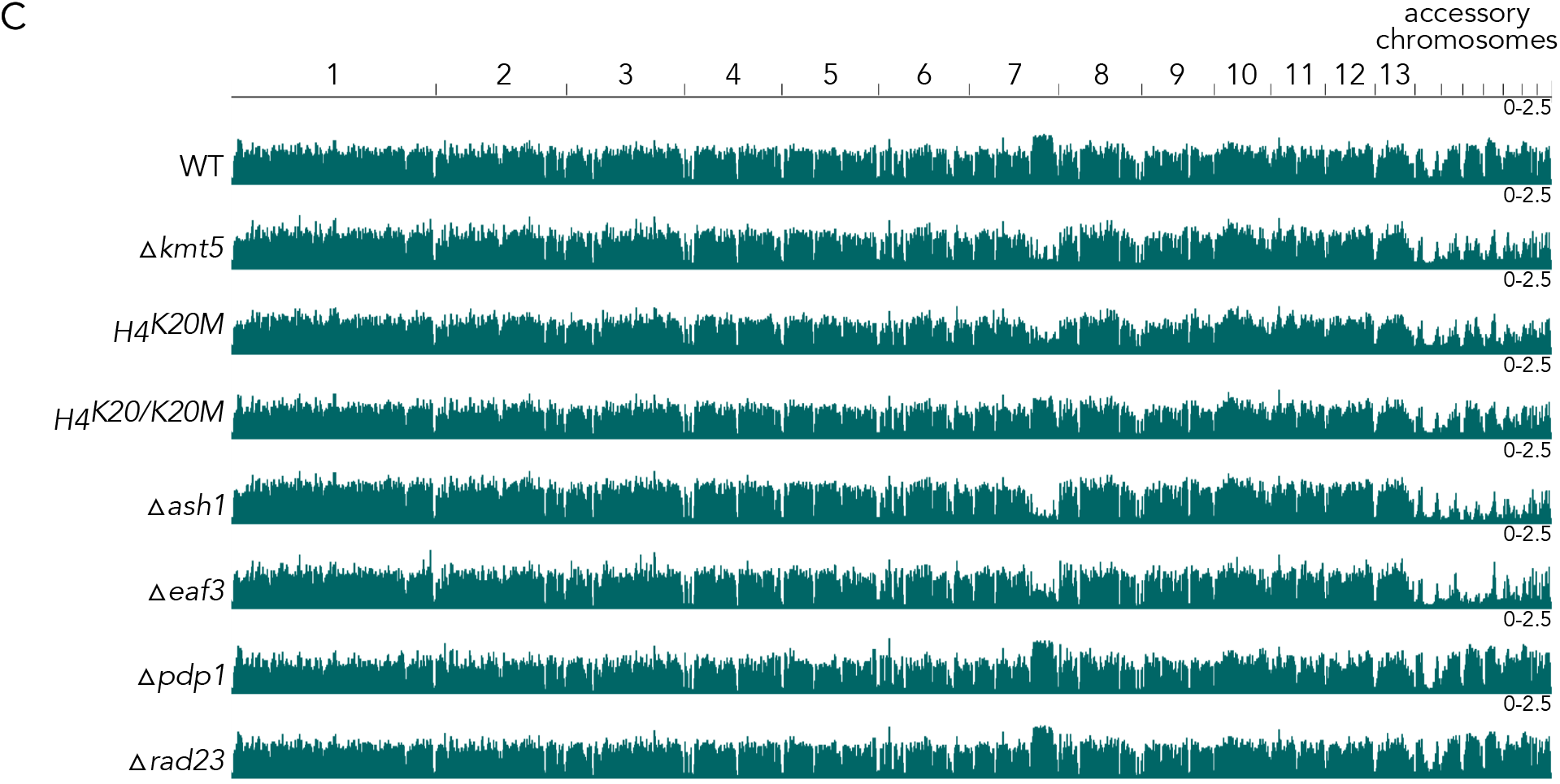
Genome-wide distribution of H4K20me3, H3K36me3 and H3K27me3 in all mutants generated in this study. There are no ChIP-seq data for H3K36me3 in the tagged Kmt5 strains.

**Figure S9.**
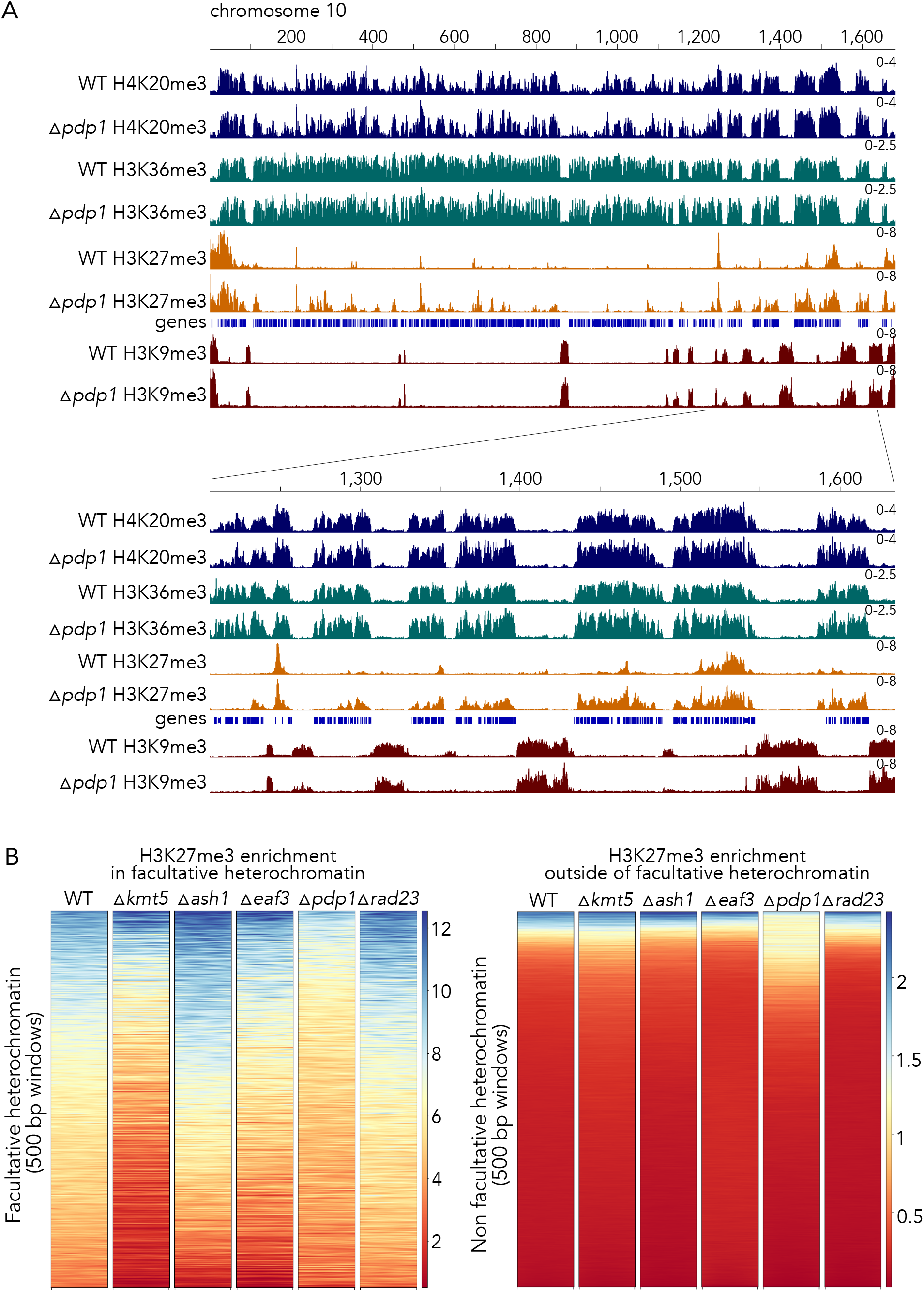
ChIP-seq of the Δ*pdp1* revealed an increase in H3K27me3 outside of facultative heterochromatin regions in WT. A) ChIP-seq tracks (chromosome 10 as an example region) show that H3K27me3 enrichment increases in gene-dense regions that are also enriched with H4K20me3 and H3K36me3 but not H3K9me3. B) H3K27me3 levels decrease in facultative heterochromatin but increase in some non-facultative heterochromatin regions.

**Figure S10.**
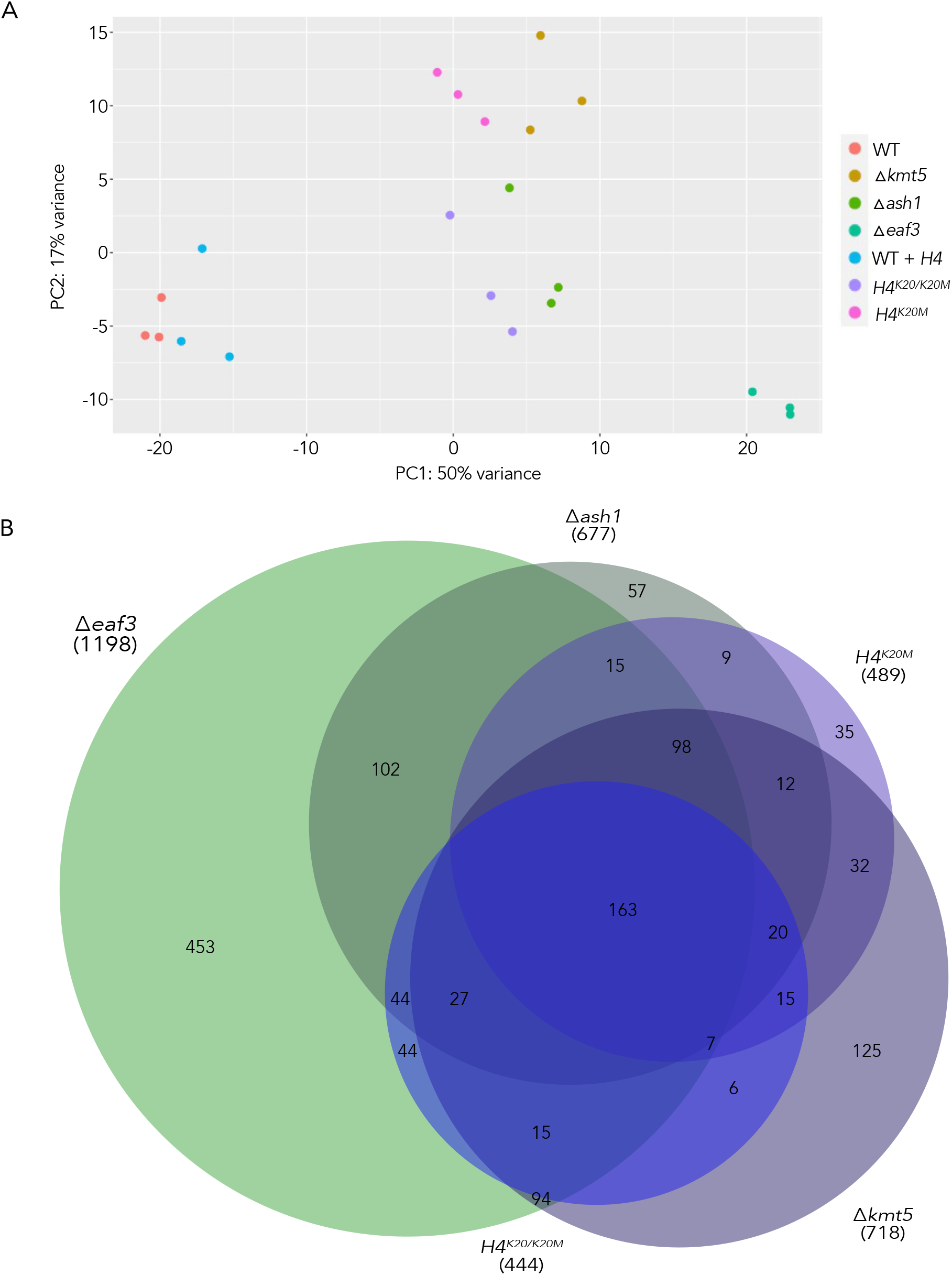
RNA sequencing of mutant strains. A) Principal Component Analysis (PCA) plot of all sequenced replicate strains. B) Venn diagram of upregulated genes in all mutants compared to wild type.

## Methods

ChIP analysis

RNA-seq analysis

Hi-C analysis

ChIP protocol

Stranded mRNA library protocol

Modified NEB Illumina library protocol

*Zymoseptoria* transformation protocol

