## Supplementary material for "H4K20me3 controls Ash1-mediated H3K36me3 and transcriptional silencing in facultative heterochromatin": Suppl_Method_IlluminaLib

Protocol to construct ChIP-sequencing libraries for Illumina sequencing
(adapted from Ferraro and Lewis, 2018)

Reagents:

- Cytiva SeraMag SpeedBeads, magnetic carboxylate modified particles (ThermoFisher, 651521050250; see Rohland and Reich, 2012).
- Double strand universal adaptor oligonucleotides for Illumina sequencing (NEB or comparable supplier)
- Dual index primers for library amplification (NEB or comparable supplier).
- NEB Ultra II end repair and dA-tailing module (NEB, E7546S)
- 10 mM Tris-HCl, pH 8
- 30 % PEG-8000 in 1.25 M NaCl
- 80 % ethanol
- T4 DNA ligase (NEB, M0202S)
- 2x Quick Ligase Buffer (QLB)
- Phusion DNA polymerase (NEB or comparable supplier)
- 5x Phusion HF buffer (NEB, B0518S)
- 20 mM dNTPs (NEB or comparable supplier)
- Qubit dsDNA HS assay kit (ThermoFisher, Q32851)

Materials:

- 0.2 mL PCR tubes (8-strip tubes work best)
- Magnetic rack for PCR tubes
- Benchtop centrifuge
- PCR machine
- Qubit fluorometer

2x Quick Ligase Buffer

| | |
|---|---|
| 132 mM | Tris-HCl pH 7.6 |
| 20 mM | $MgCl_2$ |
| 2 mM | DTT |
| 2 mM | ATP |
| 15 % | PEG 6000 |

1. Perform End Repair

Prepare reagents, thaw end repair buffer on ice.

*Mix thoroughly to make sure all buffer components are in solution.*

- mix:
  - 25.5 µL (10 µL ChIP DNA plus 15.5 ul 1/10 TE)
  - 3.0 µL 10X NEBNext Ultra II end repair reaction buffer
  - 1.5 µL NEBNext Ultra II end repair enzyme mix
  - =30 µL total volume *(mix thoroughly – pipet up and down)*

Incubate in a PCR machine:
30 min at 20°C
30 min at 65°C
Hold at 4°C

1.1 Bead clean-up for switching to ligase buffer, remove small fragments

- mix:
  - 30 µL from end-repair and tailing reaction
  - 54 µL SeraMag SpeedBeads (1.8 volumes of beads)
  - 54 µL 30% PEG-8000 in 1.25 M NaCl
- mix well by pipetting up and down or vortex
- incubate at room temperature for 5-10 min
- place on magnetic rack, wait until supernatant is clear
- remove and discard supernatant
- keep sample on magnetic rack, add 200 µL of freshly prepared 80% ethanol
- incubate for ~30 seconds
- remove and discard all supernatant
- repeat 80% ethanol wash
- quick spin, use a 20 ul pipet tip to remove ethanol, dry for ~1 min (on magnetic rack).
- remove from magnetic rack, add 22 µL 10 mM Tris-HCl, pH 8 (pre-heated to ~65°C)
- vortex, incubate beads for 5 min at room temperature, spin down briefly
- place on magnetic rack for 2 min
- transfer 20 µL to fresh PCR tube

2. Ligate annealed adapters to DNA fragments

-mix (total volume 50 μL):

| | |
|---|---|
| 20 | μL 'A'-tailed DNA |
| 25 | μL 2x Quick Ligase Buffer (homemade, see below) |
| 2.5 | μL T4 DNA ligase (400 U/μL, NEB) |
| 2.5 | μL annealed adapter pair (use 0.75 uM for <10 ng ChIP DNA) |

(make master mix containing 2x QLB, T4 DNA ligase and adapter, mix well by pipetting up and down; add adapter right before aliquoting to avoid adapter self-ligation)

-mix DNA and ligation mix by pipetting up and down
-incubate for 60 minutes at room temperature or 20°C in a PCR machine (longer is fine)

2.1 Bead Clean up

Two consecutive bead clean ups - CRITICAL STEP! Removal of PEG-containing ligation buffer and (in the second clean-up) self-ligated adaptors is essential for successful library amplification

- add 60 μL SeraMag SpeedBeads (1.2X volume)
- mix well by pipetting up and down or vortex
- incubate 5-10 min at room temperature
- place on magnet, wait until supernatant is clear
- remove and discard supernatant
- add 200 μL freshly prepared 80% ethanol without removing beads from magnet
- incubate for ~30 seconds
- remove and discard all supernatant
- repeat 80% ethanol wash
- quick spin, use a 20 ul pipet tip to remove ethanol, dry for ~1 min (on magnetic rack)
- remove tubes from magnet
- add 52 μL (first clean up) 10 mM Tris-HCl, pH 8 (pre-heated to ~65°C)
- vortex, incubate beads for 5-10 min at room temperature, spin down briefly
- place tubes on magnet for 2 min
- transfer 50 μl of supernatant to fresh PCR tube
- repeat this bead clean-up procedure and **elute in 20 μL**

The first step gets rid of the viscous ligase buffer with extra PEG and ligation enhancer (crowding reagents to improve ligation efficiency), the second step removes fragments below 200 bp (i.e., adaptor dimers at ~150 bp).

### 3. Amplify by PCR

Mix (total volume 20 µL):

| | |
|---|---|
| 7.5 | µL adaptor-ligated DNA fragments |
| 5 | µL sd$H_2O$ |
| 2.5 | µL of 10 µM i7 primer + 10 µM i5 primer premixed |
| 4 | µL 5x Phusion HF buffer |
| 0.5 | µL dNTP (20 mM) |
| 0.5 | µL Phusion DNA polymerase |

PCR conditions:
- denature at 98°C for 30 sec

 **12 cycles (fewer cycles are preferred)**
 98°C for 10 seconds
 50°C for 30 seconds
 72°C for 30 seconds
- final extension: 72°C for 5 minutes
- hold at room temperature

### 4. Final bead clean-up

- add 25 µL SeraMag SpeedBeads to 20 µL PCR (1.25x volume)
- mix well by pipetting up and down or vortex
- incubate 5-10 min at room temperature
- place on magnet, wait until supernatant is clear.
- remove and discard supernatant
- add 200 µL freshly prepared 80% ethanol without removing beads from magnet
- incubate for ~30 seconds
- remove and discard all supernatant
- repeat 80% ethanol wash
- quick spin, use a 20 ul pipet tip to remove ethanol, dry for ~1 min (on magnetic rack).
- remove tubes from magnet
- add 22 µL 10 mM Tris-HCl, pH 8 (pre-heated to ~ 65°C)
- vortex, incubate beads for 5-10 min at room temperature, spin down briefly
- place tubes on magnet for 2 min
- move 20 ul of supernatant to fresh PCR tube
- run 4 µL of the library on a 1.5% agarose gel to confirm correct size distribution
- quantify DNA concentration with a Qubit fluorometer (HS DNA kit)
