## Supplementary material for "H4K20me3 controls Ash1-mediated H3K36me3 and transcriptional silencing in facultative heterochromatin": Suppl_Methods_HiC

## Hi-C analysis

## used HiCUP v0.8.3 for truncating and mapping, see modified config files

#configuration file for the hicup Perl script
########################################################################

#Directory to which output files should be written (optional parameter)
#Set to current working directory by default 
Outdir:


#Number of threads to use
Threads: 12


#Suppress progress updates (0: off, 1: on)
Quiet:0


#Retain intermediate pipeline files (0: off, 1: on)
Keep:0


#Compress outputfiles (0: off, 1: on)
Zip:1


#Path to the alignment program (Bowtie or Bowtie2)
#Remember to include the executable Bowtie/Bowtie2 filename.
#Note: ensure you specify the correct aligner i.e. Bowtie when 
#using Bowtie indices, or Bowtie2 when using Bowtie2 indices. 
#In the example below Bowtie2 is specified.
Bowtie2: /Applications/bowtie2/bowtie2


#Path to the reference genome indices
#Remember to include the basename of the genome indices
Index: ./Zt09_chromosome18 


#Path to the genome digest file produced by hicup_digester
Digest: ./Digest_Zt09_genome_DpnII_None_13-34-46_05-05-2020.txt


#FASTQ format (valid formats: 'Sanger', 'Solexa_Illumina_1.0', 'Illumina_1.3' or 'Illumina_1.5')
#If not specified, HiCUP will try to determine the format automatically by analysing
#one of the FASTQ files. All input FASTQ will assumed to be in this format
Format: 


#Maximum di-tag length (optional parameter)
Longest: 800


#Minimum di-tag length (optional parameter)
Shortest: 50


#FASTQ files to be analysed, placing paired files on adjacent lines
./sample1_R1.fastq.gz
./sample1_R2.fastq.gz

########################################################################

## used output of hicup_mapper as input for analysis in homer

makeTagDirectory sample1_HiC_homer sample1_HiC_1.sam,sample1_HiC_2.sam -tbp 1

cp -r sample1_HiC_homer/ sample1_HiC_homer_filter/

makeTagDirectory sample1_HiC_homer_filter -update -restrictionSite GATC -both -genome Zt09_chromosomes_18.fasta -removePEbg

## Figure 2, genome wide overview

analyzeHiC sample1_HiC_homer_filter -res 50000 -superRes 50000 -norm > sample1_50kb_50kb_filter_norm.txt

## Suppl figure, chromosome 7

analyzeHiC sample1_HiC_homer_filter -res 10000 -superRes 10000 -chr sample1_chr_07 -norm > sample1_10kb_10kb_filter_chr7_norm.txt
