## Supplementary material for "H4K20me3 controls Ash1-mediated H3K36me3 and transcriptional silencing in facultative heterochromatin": Suppl_Method_Transformation

### Transformation of *Zymoseptoria tritici* with *Agrobacterium tumefaciens*

**Antibiotics and reagents**
Kanamycin (50 mg/mL)
Carbenicillin (100 mg/mL)
Rifampicin (100 mg/mL)
Cefotaxime (100 mg/mL)
Timentin (150 mg/mL)
Hygromycin (100 mg/mL)
G418 (100 mg/mL)
Acetosyringone (100 mM)

**Media**
YMS
LB
Induction medium

**Material**
Sterile Hybond-N+ membranes (Biodyne B Nylon Membrane, 0.45 μM)

| YMS (Yeast Malt Sucrose) | |
|---|---|
| $dH_2O$ | to 1000 mL |
| Yeast extract | 4 g |
| Malt extract | 4 g |
| Sucrose | 4 g |
| (Agar | 16 g) |

**Autoclave**

| LB | |
|---|---|
| $dH_2O$ | to 1000 mL |
| Yeast extract | 5 g |
| Tryptone | 10 g |
| NaCl | 10 g |
| (Agar | 16 g) |

**Autoclave**

| *Agrobacterium* Induction Minimal Medium (IMM) | |
|---|---|
| $dH_2O$ | to 1000 mL |
| $K_2HPO_4$ | 2.05 g |
| $KH_2PO_4$ | 1.45 g |
| $NH_4NO_3$ | 0.5 g |
| NaCl | 0.15 g |
| $FeSO_4$ | 0.0025 g |
| $CaCl_2$ | 0.01 g |
| $MgSO_4$ | 0.25 g |
| Glucose | 0.9 g |
| $MES \cdot H_2O$ (free acid) | 5.33 g |
| Glycerol | 5 mL |
| Vogel trace elements | 20 μL |
| (Agar | 16 g) |

**Autoclave**

| Vogel trace elements medium | |
|---|---|
| $dH_2O$ | 95 mL |
| citric acid$\cdot H_2O$ | 5.0 g |
| $ZnSO_4 \cdot 7H_2O$ | 5.0 g |
| $Fe(NH_4)_2SO_4 \cdot 6H_2O$ | 1.0 g |
| $CuSO_4 \cdot 5H_2O$ | 0.25 g |
| $MnSO_4 \cdot 1H_2O$ | 0.05 g |
| $H_3BO_3$ | 0.05 g |
| $Na_2MoO_4$ | 0.05 g |

Bring volume to 100 ml and add 1 ml of chloroform (as a preservative). Store 4° C.

Approximately one week before transformation:
- Streak out the desired *Zymoseptoria* strain from –80°C glycerol stock on YMS plate and incubate at 18°C for 4-5 days (some mutants may need more time)
- Transform *Agrobacterium tumefaciens* strain AGL1 with the confirmed vector (see AGL1 transformation protocol)
- Make sure you have all required media and sterile Hybond-N+-membrane

Two days before transformation:
- Inoculate a single colony of transformed AGL1 in 2.5 mL of LB + Rifampicin, Carbenicillin and Kanamycin in a test tube. Incubate this pre-culture for approximately 12-24 hours at ~28°C at ~200 rpm.

**For 10 ml LB + antibiotics**
10mL LB medium
10 µL Carbenicillin (stock 100 mg/mL, final concentration [f.c.] 100 µg/mL)
5 µL Rifampicin (stock 100 mg/mL, f.c. 50 µg/mL)
10 µL Kanamycin (stock 50 mg/mL, f.c. 50 µg/mL)

Evening before transformation:
**Induce AGL1**
- Spin down ~1 mL of AGL1 LB pre-culture (5 min, 5,000 rpm, RT) and resuspend in 1 mL induction medium + antibiotics and acetosyringone
- Measure $OD_{590}$ of the AGL1 pre-cultures (make 1:10 dilution in water)
- Dilute AGL1 to an $OD_{590}$ of 0.15 in induction medium in a total volume of 5 mL. Use a 50 mL Falcon tube for over-night induction.
- Take 750 µL of diluted Agrobacterium culture ($OD_{590}$ 0.15) and measure $OD_{590}$ to check if the dilution is correct (target $OD_{590}$ = 0.15). Incubate the culture in a sterile 50 mL Falcon tube at ~28°C at ~200 rpm overnight.

**For 10 mL induction medium with antibiotics and acetosyringone** (IM)
10 mL induction minimal medium (IMM)
10 µL Carbenicillin (stock 100 mg/mL, f.c. 100 µg/ml=L)
5 µL Rifampicin (stock 100 mg/mL, f.c. 50 µg/mL)
10 µL Kanamycin (stock 50 mg/mL, f.c. 50 µg/mL)
20 µL Acetosyringone (stock 100 mM, f.c. 200 µM)

**For 100 mL induction medium agar with antibiotics and acetosyringone (IM)**
100 mL induction minimal medium (IMM) with agar
100 µL Carbenicillin (stock 100 mg/mL, f.c. 100 µg/mL)
50 µL Rifampicin (stock 100 mg/mL, f.c. 50 µg/mL)
100 µL Kanamycin (stock 50 mg/mL, f.c. 50 µg/mL)
200 µL Acetosyringone (stock 100 mM, f.c. 200 µM)

- Pour induction medium (IM) plates (2-3 plates per transformation)

<u>Transformation day:</u>

**Prepare *Zymoseptoria***
- scrape off some *Z. tritici* cells from YMS plate and resuspend in 1 mL of sterile $dH_2O$
- make a 1:10 dilution of resuspended cells and measure $OD_{590}$
- prepare cell suspension with $OD_{590}$=1.5 in induction medium with antibiotics and acetosyringone (IM); 125 µL *Zymoseptoria* cell suspension per transformation plate

**Prepare AGL1**
- measure $OD_{590}$ of induced overnight AGL1 culture (should be between 0.35-0.7)
- if necessary, dilute AGL1 culture to $OD_{590}$=0.45 in liquid IM (125 µL AGL1 cell suspension per transformation plate); if culture has not reached $OD_{590}$=0.45, an $OD_{590}$ >0.35 is fine)

- mix *Zymoseptoria* ($OD_{590}$=1.5) and AGL1 ($OD_{590}$=0.45) at a 1:1 ratio in 1.7 mL tube (250 µL of mixture per transformation plate), incubate for ~10 min

- during incubation, place sterile Hybond-N+ membranes on induction medium plates with tweezers

- plate 250 µL of *Zymoseptoria*/AGL1 mixture on membrane

- incubate plates (sealed with parafilm) at room temperature (~20-22°C) for two days

<u>Two days after transformation:</u>

- transfer Hybond-N+ membrane to selection plate; at this stage colonies are not visible
- plates need to contain antibiotics to select for fungal transformants and to select against AGL1

- incubate plates (sealed with parafilm) at 18°C until colonies appear, usually after 7-14 days (depending on strain, selection, and possible growth defects; also pick small and slow growing colonies)

- pick colonies (make sure to pick single colonies) from transformation plate onto YMS selection plates (still fungal and bacterial selection) and screen for correct integration by PCR (quick DNA extraction and screening protocol)

For 100 mL YMS agar
100 mL YMS agar
100 µL Timentin (stock 150 mg/mL, f.c. 150 µg/mL)
200 µL Cefotaxime (stock 100 mg/mL, f.c. 200 mg/mL)

Depending on selection marker:
100 µL Hygromycin (stock 100 mg/mL, f.c. 100 µg/mL)
100 µL G418 (stock 100 mg/mL, f.c. 100 µg/mL)
