## Supplementary material for "H4K20me3 controls Ash1-mediated H3K36me3 and transcriptional silencing in facultative heterochromatin": Suppl_Method_RNALib

Strand-specific RNA sequencing library preparation
(adapted from Zhong et al., 2011)

Reagents:

- oligo d(T)25 magnetic beads (NEB, S1419S)
- SeraMag SpeedBeads, magnetic carboxylate modified particles (Thermo Scientific, 651521050250); see Rohland and Reich, 2012.
- SuperScript III reverse transcriptase and 5X SuperScript III first-strand buffer (Thermo Scientific, 18080044)
- nuclease-free or DEPC-treated $H_2O$
- random hexamers $(N)_6$ (IDT)
- RNasin Plus ribonuclease inhibitor (Promega, N2611)
- Actinomycin D (Adipogen, BVT-0089-M005)
- dNTP solution set (dATP, dCTP, dGTP, dTTP; NEB, N0446S)
- dUTP solution (NEB, N0459S)
- 10X NEBuffer 2 or 10X NEBuffer 2.1 (NEB, B7002S)
- RNase H (NEB, M0297L)
- DNA polymerase I (NEB, M0209S)
- Illumina iTruSeq adapter and i7 and i5 primers (NEB or comparable supplier)
- NEB Ultra II end repair and dA-tailing module (NEB, E7546S)
- T4 DNA ligase (NEB, M0202L)
- Uracil DNA glycosylase (UDG; NEB, M0280S)
- Phusion polymerase and 10X Phusion reaction buffer (NEB or comparable supplier)

Solutions:

*Note: All solutions should be prepared with nuclease-free or DEPC-treated sterile $H_2O$ when possible.*

- 2X binding buffer
  - 1 M LiCl
  - 20 mM Tris-HCl (pH 7.5)
  - 2 mM EDTA (pH 8.0)
  - 1% lithium dodecyl sulfate
- wash buffer
  - 0.15 M NaCl
  - 10 mM Tris-HCl (pH 7.5)
  - 1 mM EDTA (pH 8.0)
- 1X TE buffer
  - 10 mM Tris-HCl (pH 7.5)
  - 1 mM EDTA (pH 8.0)
- 2X Quick Ligase Buffer (QLB; see updated ChIP Illumina library protocol for dual indexing for additional information)
  - 132 mM Tris-HCl (pH 7.6)
  - 20 mM $MgCl_2$

  - 2 mM DTT
  - 2 mM ATP
  - 15% polyethylene glycol, molecular weight 6000 (PEG-6000)
- 80% ethanol (freshly prepared with DEPC-treated $H_2O$)
- 100 mM DTT

Methods:
PolyA RNA isolation and fragmentation

1. Prepare oligo d(T)25 beads:
   a. Equilibrate beads to room temperature and mix thoroughly,
   b. Transfer appropriate volume of bead slurry (20 µl per sample) to a 1.7 ml microcentrifuge tube,
   c. Place tube containing bead slurry on magnetic rack to collect beads,
   d. Wash beads twice with equal volume of 1X binding buffer. *Note: 2X binding buffer should be diluted with 1X TE buffer.*

2. Resuspend oligo d(T)25 beads in appropriate volume of 2X binding buffer (50 µl per sample).

3. Add 50 µl oligo d(T)25 beads to PCR tubes containing 50 µl total RNA (in $H_2O$) and mix well by pipetting.

4. Heat samples to 65°C for 2 min in a thermocycler with heated lid to denature the RNA and facilitate mRNA binding.

5. Incubate samples at room temperature for 5-10 min with occasional gentle mixing.

6. Place samples on magnetic rack to collect beads. Without disturbing bead pellet, carefully remove and discard cleared supernatant.

7. Remove samples from the magnetic rack. To remove unbound RNA, add 150 µl wash buffer and mix well by pipetting.

8. Place samples on magnetic rack to collect beads. Without disturbing bead pellet, carefully remove and discard cleared supernatant.

9. Repeat washing (steps 7-8).

10. Remove samples from magnetic rack. Add 50 µl 1X TE buffer and mix well by pipetting.

11. Incubate samples at 80°C for 2 min to elute mRNA. Immediately place samples on ice for 1 min.

12. Add 50 µl 2X binding buffer and mix well by pipetting.

13. Repeat binding and washing (steps 4-9).

14. Remove samples from the magnetic rack. Add 30 µl ice-cold 1X superscript III first-strand buffer and mix well by pipetting. *Note: Lithium ions inhibit reverse transcriptase. This step is critical to remove lithium ions, salts, and detergent from samples.*

15. Place samples on magnetic rack to collect beads. Without disturbing bead pellet, carefully remove and discard cleared supernatant. Carefully inspect samples and remove remaining buffer with a 10 µl pipet.

16. Remove samples from magnetic rack. Resuspend bead-bound mRNA in 12 µl of 2X superscript III first-strand buffer supplemented with 10 mM DTT.

17. Incubate samples at 94°C for exactly 3 min to fragment mRNA. Immediately place samples on ice. *Note: Fragmentation time can be optimized to generate libraries of different insert sizes (see Zhong et al., 2011 for additional information).*

18. Place samples on magnetic rack for 5 min to collect beads. Transfer cleared supernatant (10 µl) containing fragmented mRNA to a fresh PCR tube. *Note: Fragmented mRNA can be stored at -80°C.*

First-strand cDNA synthesis

19. Prepare the following mixture on ice:

| Reagent | Volume (µl) |
|---|---|
| Fragmented mRNA (in 2X SuperScript III first-strand buffer) | 10 |
| Random hexamers $(N)_6$ (1 µg/µl) | 0.5 |
| RNase inhibitor (40 U/µl) | 0.5 |
| Total | 11 |

20. Heat samples at 50°C for 1 min. Immediately place samples on ice.

21. Prepare the following reverse transcription (RT) reaction mixture on ice:

| Reagent | Volume (µl) |
|---|---|
| Nuclease-free $H_2O$ | 6.75 |
| Actinomycin D (1 µg/µl) | 0.12 |
| DTT (100 mM) | 1 |

| | |
|---|---|
| dNTP mix (20 mM) | 0.63 |
| Superscript III reverse transcriptase | 0.5 |
| Total | 9 |

22. Add 9 µl of the RT reaction mixture to each sample (11 µl). Mix well by pipetting.

23. Heat samples to 25°C for 10 min followed by 50°C for 50 min in a thermocycler with heated lid.

24. Immediately purify RT reactions using 1.8 volumes (36 µl) SeraMag SpeedBeads. Mix well by pipetting and incubate samples on ice for 15 min.

25. Place samples on magnetic rack for 5 min to collect beads and proceed with purification as described below in *General method for nucleic acid purification with SeraMag SpeedBeads*.

26. Elute RNA/cDNA hybrid with 12 µl nuclease-free $H_2O$.

### Second-strand cDNA synthesis (with dUTP)

27. Prepare the following second strand reaction mixture on ice:

| Reagent | Volume (µl) |
|---|---|
| Nuclease-free $H_2O$ | 1.3 |
| NEBuffer 2 | 1.5 |
| dNTP mix (10 mM; dATP, dCTP, dGTP, and dUTP) | 1 |
| RNase H (5 U/µl) | 0.2 |
| DNA polymerase I (10 U/µl) | 1 |
| Total | 5 |

28. Add 5 µl of the second strand reaction mixture to each RNA/cDNA sample (10 µl). Mix well by pipetting and incubate at 16°C for 2.5 h.

29. Purify double-stranded DNA (dsDNA) using 1.8 volumes (27 µl) SeraMag SpeedBeads as described below in *General method for nucleic acid purification with SeraMag SpeedBeads*.

30. Elute dsDNA with 22 µl 1X TE buffer preheated to 50°C. *Note: Purified dsDNA can be stored at -20°C.*

### End-repair, dA-tailing, and adaptor ligation

31. Perform end-repair, A-tailing, and adaptor ligation as described in the updated ChIP Illumina library protocol (Step 1) using the annealed universal adapter pair (5 µM). *Note: Final volume of the adaptor-ligated library is 50 µl.*

### Buffer exchange and double-sided size selection

32. Purify library using 1 volume (50 µl) SeraMag SpeedBeads as described below in *General method for nucleic acid purification with SeraMag SpeedBeads*.

33. Elute library with 52 µl 1X TE buffer preheated to 50°C.

34. Perform right-side size selection (removal of large fragments) by adding exactly 0.5 volumes (25 µl) SeraMag SpeedBeads to each library. Mix well by pipetting and incubate at room temperature for 5 min. Proceed with Step 35 during incubation. *Note: Proper size selection is a critical step (see Zhong et al., 2011 for additional information).*

35. Prepare 2X SeraMag SpeedBeads:
    a. Ensure beads are equilibrated to room temperature and mix thoroughly,
    b. Transfer twice the appropriate volume of bead slurry (14 µl per sample) to a 1.7 ml microcentrifuge tube,
    c. Place tube containing bead slurry on magnetic rack to collect beads,
    d. Carefully remove half the volume of cleared supernatant without disturbing bead pellet,
    e. Remove tube from magnetic rack and mix thoroughly to resuspend beads.

36. Place samples on magnetic rack for 5 min to collect beads. Without disturbing bead pellet, carefully transfer cleared supernatant (72 µl) to a fresh PCR tube containing 0.3 volumes (13.9 µl) 2X SeraMag SpeedBeads for left-side size selection (removal of small fragments). *Note: This is equivalent to 0.8 volumes considering the volume of beads used in Step 34 (see Illumina technical bulletin* "Double-sided size selection and bead clean-up" *for additional information).*

37. Mix well by pipetting and incubate at room temperature for 5 min. Proceed with purification as described below in *General method for nucleic acid purification with SeraMag SpeedBeads*.

38. Elute library with 22 µl 1X TE buffer preheated to 50°C. *Note: Purified, size-selected library can be stored at -20°C.*

### UDG digest and PCR enrichment

39. Digest the second strand DNA by adding 0.5 µl UDG to 10 µl size-selected library. Incubate digestions at 37°C for 15 min.

**40.** Prepare the following PCR mixture on ice:

| Reagent | Volume (µl) |
|---|---|
| Nuclease-free $H_2O$ | 1.5 |
| 10X Phusion reaction buffer | 2 |
| dNTP mix (20 mM) | 0.5 |
| Premixed i7 primer and i5 primer (10 µM) | 5 |
| Phusion polymerase | 1 |
| UDG-digested, size-selected library | 10 |
| Total | 20 |

**41.** Perform initial denaturation at 98°C for 30 s, followed by 8-14 cycles of amplification (98°C for 10 s, 50°C for 30 s, and 72°C for 30 s) and hold at room temperature. *Note: Use 8-10 cycles if starting with >5 ug total RNA, 10-12 cycles if starting with 1-5 ug total RNA, or 14 cycles if starting with 0.5-1 ug total RNA.*

Purify library using 1.4 volumes (28 µl) SeraMag SpeedBeads as described below in *General method for nucleic acid purification with SeraMag SpeedBeads.*

**42.** Elute library with 22 µl 1X TE buffer (pH 8.0) preheated to 50 °C. *Note: Purified, enriched library can be stored at -20 °C.*

**Additional methods:**

**General method for nucleic acid purification with SeraMag SpeedBeads**

**1.** Add indicated amount of SeraMag SpeedBeads to sample.

**2.** Mix well by pipetting up and down at minimum of 10 times.

**3.** Incubate samples at room temperature for 5 min. *Note: Unless specified in the protocol, prolonged incubation or incubation at low temperature will increase binding of small DNA fragments (e.g., adapter dimers).*

**4.** Place samples on magnetic rack for 5 min (or until solution is clear) to collect beads. *Note: Keep samples on magnetic rack for steps 4-8.*

**5.** Carefully remove cleared supernatant without disturbing bead pellet.

**6.** Add 180 µl 80% ethanol and incubate for 30 s. Carefully remove ethanol wash without disturbing bead pellet.

7. Repeat 80% ethanol wash once. Carefully inspect samples and remove remaining ethanol with a 10 µl pipet.

8. Air dry samples for 5 min with lid open. Do not overdry samples. *Note: Samples are sufficiently dry when small cracks begin to appear on bead or bead ring.*

9. Remove samples from magnetic rack. Add indicated volume of appropriate solution for elution and mix well by pipetting.

10. Incubate mixture at room temperature for 2-5 min.

11. Place samples on magnetic rack for 2 min (or until solution is clear) to collect beads.

12. Transfer cleared supernatant (volume of solution added for elution less 2 µl) to a fresh PCR tube.
