## Supplementary material for "H4K20me3 controls Ash1-mediated H3K36me3 and transcriptional silencing in facultative heterochromatin": Suppl_Method_ChIPZymo

### ChIP (chromatin immunoprecipitation) protocol for *Zymoseptoria tritici*

#### Buffers and reagents

**ChIP lysis buffer**
50 mM HEPES-NaOH, pH 7.5
**90 mM NaCl**
1 mM Na-EDTA, pH 8.0
1% Triton X-100
0.1% DOC
+ proteinase inhibitors (PMSF, Leupeptin, Pepstatin, E-64)

**10 mL of ChIP lysis buffer**
500 µL of 1 M HEPES-NaOH, pH 7.5
**300 µL of 3 M NaCl**
20 µL of 0.5 M EDTA, pH 8
1 mL of 10% Triton X-100
100 µL of 10% DOC (Na-Deoxycholate)
**100 µL of 0.1 M PMSF**
**10 µL of 1000x Leupeptin**
**10 µL of 1000x E-64**
**30 µL of 333x Pepstatin**
Fill-up with $sdH_2O$ to 10 mL
Store at 4°C.

**ChIP lysis buffer plus 0.5 M NaCl**
50 mM HEPES-NaOH, pH 7.5
**500 mM NaCl**
1 mM Na-EDTA, pH 8
1 % Triton X-100
0.1 % DOC

**20 mL of ChIP lysis buffer plus 0.5M NaCl**
1 mL of 1 M HEPES-NaOH, pH 7.5
**3.33 µL of 3 M NaCl**
40 µL of 0.5 M EDTA, pH 8
2 mL of 10% Triton X-100
200 µL of 10% DOC (Na-Deoxycholate)
Fill-up with $sdH_2O$ to 20 mL
Store at 4°C.

**ChIP LiCl wash buffer**
10 mM Tris-HCl, pH 8.0
250 mM LiCl
0.5 % IGEPAL CA-630
0.5 % DOC
1 mM Na-EDTA, pH 8

**20 mL of ChIP LiCl wash buffer**
200 µL of 1M Tris-HCl, pH 8.0
1 mL of 5 M LiCl
100 µL of 100 % IGEPAL
1 mL of 10% DOC (Na-Deoxycholate)
40 µL of 0.5 M EDTA, pH 8
Fill-up with $sdH_2O$ to 20 mL
Store at 4°C.

20 % Formaldehyde (methanol-free)
1 M $CaCl_2$
0.5 M EGTA
magnetic beads (ThermoFisher, Dynabeads Protein A or Protein G; 10001D, 10003D)
antibodies
1 X PBS
2.5 M glycine
1 X TE
ChIP DNA Clean & Concentrator (Zymo Research D5205)

**50 mL TES**
2.5 mL of Tris-HCl, pH 8.0
1 mL of 0.5 M EDTA, pH 8
5 mL 10 % SDS
41.5 mL of $sdH_2O$
Store at RT.

ChIP buffers (without proteinase inhibitors) can be stored at 4°C for at least three months.

### ChIP protocol

#### Preparation of *Zymoseptoria* cells for *in vitro* ChIP

Grow *Zymoseptoria* strains on YMS plates (4 g yeast extract, 4 g malt extract, 4 g sucrose, 16 g agar in 1 liter water) at 18°C. Streak cells out from glycerol stock, let cells grow for 3-4 days, transfer to new plates (2-3 plates per strain) and grow for another 3-4 days.

• Scrape cells off the plate using a pipet tip and resuspend in 5 mL 1x PBS (room temperature) in 50 mL tube.
• Add 125 µL of 20 % formaldehyde and crosslink for 15 min at RT.
• Add 100 µL of 2.5 M glycine to quench formaldehyde and incubate for 5 min at RT.
• Centrifuge for 5 min, 4,000 rpm, at 4 °C.
• Remove supernatant and wash pellet with 5 mL cold 1x PBS.
• Centrifuge for 5 min, 4,000 rpm, at 4 °C.
• Remove supernatant, keep cells on ice.

→ Safe stopping point! Crosslinked cells can be snap-frozen in liquid nitrogen and stored at -80°C for a few days, possibly longer.

• Grind cells in liquid nitrogen until you have completely white, fine powder.
• Transfer ~100-300 mg of ground cells to pre-cooled (in liquid nitrogen) 1.7 mL screw cap tubes.
• Keep tubes with ground cells in liquid nitrogen until adding the ChIP lysis buffer (including proteinase inhibitors!) or until storing them at -80°C.

→ Safe stopping point! Ground cells can be stored at -80°C (for at least 1 month).

#### ChIP Day 1

• Add ChIP lysis buffer (add proteinase inhibitors before use) to the ground cells, at a ratio of 5 µL of ChIP lysis buffer per 1 mg cells (maximum of 1 mL or MNase digestion must be adjusted). Vortex to mix cells and lysis buffer.
(when more than 1 mL of chromatin is required, use multiple tubes per replicate and pool the resuspended ground cells in an ice-cold 15 mL tube, then transfer equal volumes (max. 1 mL) to separate tubes).
• Incubate for 10-15 min on ice.
• Add 2 µL of $CaCl_2$ to each tube.
• Add 5 µL of MNase (micrococcal nuclease; NEB, M0247S, 10,000 Units).
• Incubate for 10-15 min at 37°C (incubator or water bath), mix every 2 min by inversion.
• Add 20 µL of 0.5 M EGTA to stop MNase digestion (on ice).
• Centrifuge for 10 min at 6,000 rpm at 4°C (RT is fine).
• Transfer the supernatant to a new tube (keep tubes on ice).
• *Optional (to increase chromatin yield):* Leave ~ 100 µL of supernatant in tube. Resuspend cell pellet in remaining supernatant, centrifuge for 5 min at 6,000 rpm at 4°C (RT is fine) and add supernatant to the tube containing the supernatant from the previous step.

• Split the samples into 125 µL and ~900 µL fractions.

  o The 125 µL sample is used as a control for the MNase digestion (or as input).
  o To the 125 µL sample add 125 µL TES (prewarmed to 65°C) and 1 µL of Proteinase K (20mg/ml).
  o De-crosslink for 6-16h at 65°C (can be done over-night or on the same day).
  o Add 250 µL $sdH_2O$ and 1.9 µL RNAse A (20 mg/mL) to the de-crosslinked sample.
  o Incubate for 2 h at 50°C.
  o Use the ChIP DNA Clean & Concentrator kit to extract DNA (use 1 mL of binding buffer). Elute in 20 µL. Store at -20°C.
  o Run 10 µL DNA on a ~1.5% TAE agarose gel, successful MNase digestion is indicated by a bright band at ~150 bp and a faint nucleosome ladder (bands at ~300 bp, ~450 bp, ~600 bp, etc.).

• To the 900 µL fraction, add 25 µL of pre-washed (see below) magnetic beads (Protein A or Protein G, depending on antibodies used) and incubate for 1-3 h at 4°C on a rotating platform.

**Pre-wash magnetic beads in ChIP lysis buffer**

o Add 10X volume of ChIP lysis buffer (including proteinase inhibitors) to the volume of magnetic beads you need for your experiment.
o Place tube in magnetic rack and wait until the magnetic beads magnetize to tube wall.
o Remove ChIP lysis buffer and repeat washing once.
o Resuspend magnetic beads in original volume in ChIP lysis buffer.

• Place tubes on magnetic rack (~ 1 min) and transfer pre-cleared lysate to a new tube
• Split pre-cleared lysate into 250 µL (at least 250 µL, more is fine) aliquots and add 3 µL of desired antibody to the lysate. *Note: titer of antibodies should be tested for new batches. In our studies between 2-4 µL of commercial antibodies performed well.*
• Incubate lysate and antibody over night at 4°C on a rotating platform.

#### ChIP Day 2

**All buffers should be ice-cold.**

• Add 25 µL of pre-washed magnetic beads (Protein A or G, see above) to the samples incubated with the antibody.
• Incubate for 1-2 h at 4°C on a rotating platform.
• Place tubes on magnetic rack (~ 1 min) and discard the supernatant.
• Add 1 mL of ChIP lysis buffer (without proteinase inhibitors) and incubate for 5 min at RT on a rotating platform.
• Magnetize the beads and discard supernatant, repeat washing with ChIP lysis buffer once more.
• Wash once with 1 mL of ChIP lysis buffer + 0.5 M NaCl as described before at RT.
• Wash once with 1 mL of LiCl wash buffer as described before at RT.
• Wash once with 1 mL of 1X TE buffer as described before at RT.
• Remove supernatant completely.
• Add 62.5 µL of pre-warmed (65°C) TES to the magnetic beads (resuspend well) and incubate for 10 min at 65°C to elute the DNA from the beads. Vortex the tubes several times during incubation.
• Place tubes on magnetic rack (~ 1 min) and save the supernatant in a new tube.
• Add another 62.5 µL of pre-warmed (65°C) TES to the magnetic beads and incubate for 10 min at 65°C to elute the remaining DNA from the beads. Vortex the tubes several times during incubation.
• Place tubes on magnetic rack (~ 1 min) and pool supernatants from bead elutions.

• De-crosslink the sample (125 µL total volume) at 65°C for 6-16 hours (overnight). Ensure that snap cap tubes cannot pop open during this time or use screw cap tubes.

#### ChIP Day 3

• Add 125 µL of sd$H_2O$ and 1.9 µL of RNAse A (20 mg/mL) to the de-crosslinked sample and incubate for 2 h at 50°C.
• Add 10 µL of Proteinase K (20 mg/mL) to the sample and incubate for 2 h at 50°C.
• Use the ChIP DNA Clean & Concentrator kit to extract DNA. Use 1 mL of binding buffer. Elute in 20 µL elution buffer.
• Store samples at -20°C.
